# GFI1 cooperates with IKAROS/IKZF1 to activate gene expression in T-cell acute lymphoblastic leukemia

**DOI:** 10.1101/2021.03.08.434336

**Authors:** Wenxiang Sun, Jingtao Guo, David McClellan, Alexandra Poeschla, Diana Bareyan, Mattie J. Casey, Bradley R. Cairns, Dean Tantin, Michael E. Engel

## Abstract

Growth factor independence-1 (GFI1) is a transcriptional repressor and master regulator of normal and malignant hematopoiesis. Repression by GFI1 is attributable to recruitment of LSD1-containing protein complexes via its SNAG domain. However, the full complement of GFI1 partners in transcriptional control is not known. We show that in T-ALL cells, GFI1 and IKAROS are transcriptional partners that co-occupy regulatory regions of hallmark T cell development genes. Transcriptional profiling reveals a subset of genes directly transactivated through the GFI1—IKAROS partnership. Among these is *NOTCH3*, a key factor in T-ALL pathogenesis. Surprisingly, *NOTCH3* transactivation by GFI1 and IKAROS requires the GFI1 SNAG domain but occurs independent of SNAG—LSD1 binding. GFI1 variants deficient in LSD1 binding fail to transactivate *NOTCH3*, but conversely, small molecules that disrupt the SNAG—LSD1 interaction while leaving the SNAG primary structure intact stimulate *NOTCH3* expression. These results identify a non-canonical transcriptional control mechanism in T-ALL which supports GFI1-mediated transactivation in partnership with IKAROS and suggest competition between LSD1-containing repressive complexes and others favoring transactivation.

## Introduction

Growth factor independence-1 (GFI1) is a zinc finger transcription factor which plays essential roles in normal and malignant myeloid and lymphoid hematopoiesis (Hock, Hamblen et al., 2004, Zeng, Yucel et al., 2004). Germline *GFI1* mutations cause severe congenital neutropenia (Person, Li et al., 2003), while *Gfi1* null mice show impaired T cell and neutrophil differentiation (Hock, Hamblen et al., 2003, Karsunky, Zeng et al., 2002, Yucel, Karsunky et al., 2003). In acute myelogenous leukemia (AML), *GFI1* mRNA expression can be used to stratify patient survival, while *GFI1* displays a dose-dependent impact on the pace of leukemic progression brought on by onco-fusion proteins, MLL-AF9 and NUP98-HOXD13 (Hones, Botezatu et al., 2016, Volpe, Walton et al., 2017). Notably, a GFI1 variant, GFI136N, generated from a single nucleotide polymorphism expressed in 3-7% of the Caucasian population, is disproportionately observed in AML patients and increases risk for AML development by 60% relative to the more common GFI136S variant (Khandanpour, Krongold et al., 2012). G*FI1* mRNA is also elevated in samples from patients with early T cell precursor acute lymphoblastic leukemia (ETP-ALL) who display a positive NOTCH signature (Khandanpour, Phelan et al., 2013). ETP-ALL is a high-risk subgroup of T-ALL (Coustan-Smith, Mullighan et al., 2009, Zhang, Ding et al., 2012), which itself is an aggressive form of acute leukemia characterized by the expansion of immature lymphoid precursor cells (Terwilliger & Abdul-Hay, 2017). The precise role of GFI1 in T-ALL is not clear.

T-ALL has a high incidence of relapse, and survival following disease recurrence is dismal. Abberant activation of NOTCH signaling is a unifying theme in T-ALL, and arises either from mutations in NOTCH receptors or NOTCH regulators. Normally, in response to ligand binding, NOTCH receptors are cleaved by *γ*-secretase to liberate their intracellular domains (NICD). NICD then partners with nuclear factors to direct the expression of NOTCH target genes. *NOTCH1*-activating mutations are found in approximately 60% of T-ALL cases (Ferrando, 2010). While γ-secretase inhibitors (GSIs) have shown anti-leukemic activity *in vitro* and in murine models, they have not been integrated into T-ALL treatment protocols because of dose limiting gastrointestinal toxicity and poor anti-leukemic efficacy (Golde, Koo et al., 2013).

Like NOTCH1, NOTCH3 promotes T cell lineage specification and leukemogenesis. *NOTCH3*-activating mutations have been identified in approximately 5% of T-ALL cases and NOTCH3 blocking antibodies exhibit potent anti-leukemic effect in T-ALL (Bellavia, Campese et al., 2000, Bellavia, Campese et al., 2002, Bernasconi-Elias, Hu et al., 2016, Waegemans, Van de Walle et al., 2014, Xu, Choi et al., 2015). Moreover, abnormal expression and/or activation of *NOTCH3* is seen in T-ALL patient samples lacking *NOTCH1*-activating mutations, reinforcing NOTCH signaling as critical for T-ALL pathogenesis and suggesting *NOTCH3* and factors controlling its expression could represent alternative therapeutic targets in this disease (Choi, Severson et al., 2017, Tottone, Zhdanovskaya et al., 2019). The development of new therapeutic strategies for T-ALL depends upon deeper understanding of its molecular underpinnings.

Here, we identify IKAROS as a frequent DNA binding partner for GFI1. GFI1 and IKAROS do not interact in classical co-immunoprecipitation (co-IP) assays, but their proximity relationship is impaired by the N383S mutation that impairs GFI1 DNA binding and the N159A mutation that impairs IKAROS DNA binding. In contrast, their interaction is not affected by LSD1 binding-deficient GFI1 variants including GFI1-P2A, -K8L or -*Δ*SNAG. We identify a strong, genome-wide correlation between GFI1- and IKAROS-regulated genes through ChIP-Seq. Genes co-occupied by GFI1 and IKAROS encode hallmark T cell development proteins such as NOTCH3, CD3, GFI1 itself, c-MYC, C-MYB and HES1. Gene expression profiling by RNA-Seq identifies a cluster of genes activated by ectopic GFI1 expression and repressed with IKAROS knockout. Interestingly, these genes include the direct GFI1/IKAROS target *NOTCH3,* a critical oncogenic factor in T-ALL. Using both CCRF-CEM and SUP-T1 T-ALL cells, we show that inducible expression of either GFI1 or IKAROS elevates NOTCH3 cell surface expression, while acute degradation of IKAROS through an IKAROS inhibitor or CRISPR/Cas9-mediated IKAROS knockout significantly attenuates GFI1-mediated NOTCH3 induction. Increased NOTCH3 cell surface expression depends upon SNAG domain amino acids that enable interaction with LSD1. Yet, LSD1 inhibition and disruption of the SNAG—LSD1 interaction augments *NOTCH3* expression, suggesting that LSD1 competes with activators of NOTCH3 expression mediated by the GFI SNAG domain. Together, these results identify a noncanonical transativation mechanism for GFI1, working in partnership with IKAROS to promote expression of *NOTCH3* and related T cell development genes, and providing new insights for therapeutic targeting in T-ALL.

## Results

### GFI1 proximitome proteomics in T-ALL cells

To identify potential GFI1 cooperating proteins in T-ALL, we applied the BioID (<u>Bio</u>tin <u>ID</u>entification) proximity-dependent labeling method to screen for vicinal proteins (Figure 1A). We generated doxycycline-inducible, GFI1-BirA*-expressing CCRF-CEM cells. In parallel, cells transduced with empty vector or BirA* only were generated as controls (Figure 1B). Cells were incubated in biotin-containing medium, treated with doxycycline and lysates prepared. Biotinylated proteins were collected with streptavidin Sepharose beads and surveyed for known GFI1 interacting partners. As expected, we detected comparable biotinylation of GFI1-BirA* and BirA*, suggesting the presence of GFI1 protein structure does not interfere with the formation of reactive biotin-AMP by BirA* in the fusion protein. Moreover, we find an altered pattern of biotinylated proteins when BirA* activity is tethered to GFI1, and enrichment for known GFI1 interacting partners, LSD1 and CoREST among biotinylated proteins in cells expressing GFI1-BirA* vs. BirA* control (Figure 1C). These data validate the technique for detecting GFI1 proximity partners proteome-wide.

**Figure 1.**
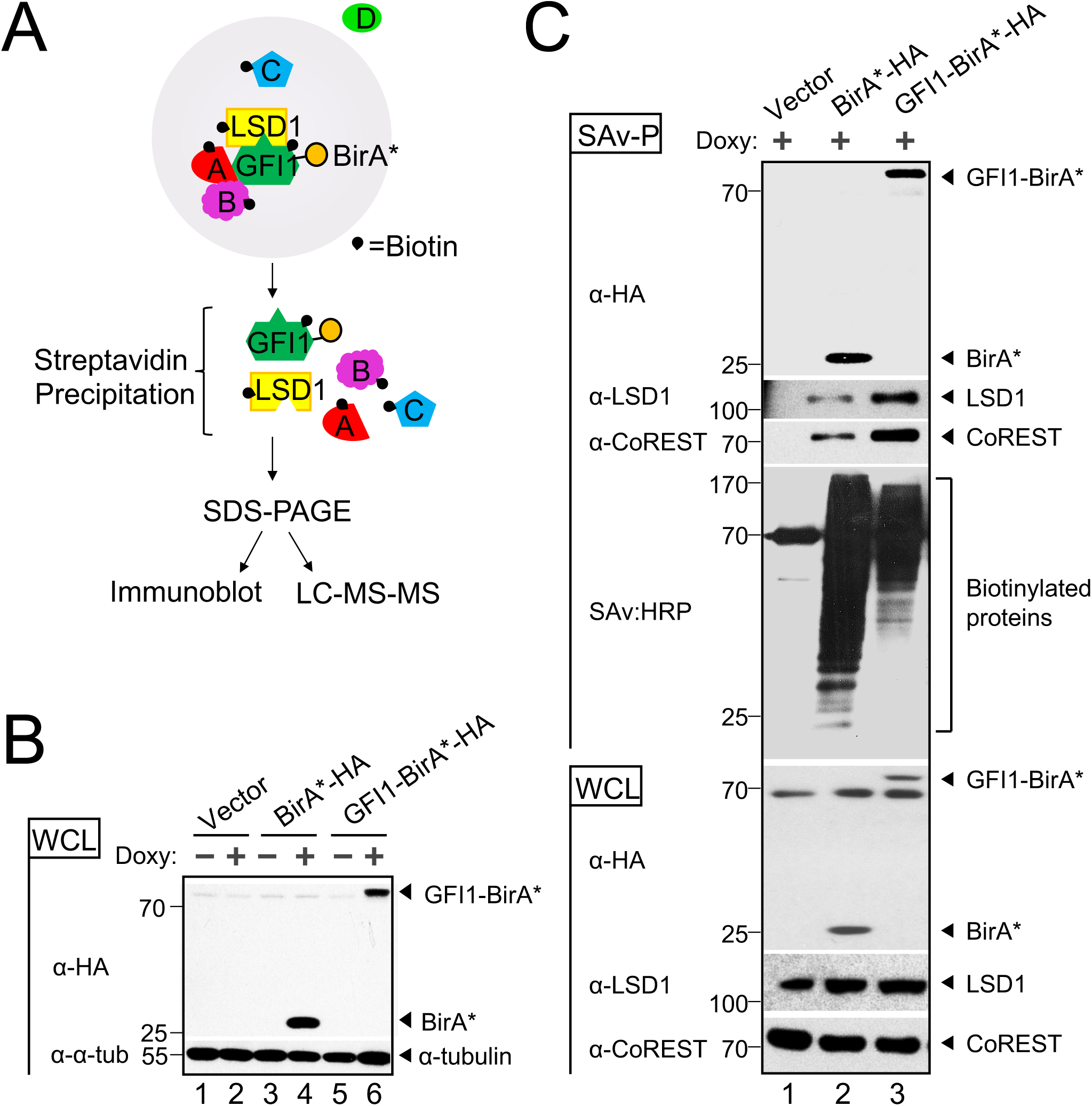
BioID proximity labeling identifies GFI1-interacting proteins. (**A**) Scheme for BioID proximity labeling method to identify GFI1-proximate proteins. Gray circle reflects the radius for biotinylation of nearby molecules such as GFI1 itself and its known binding partner LSD1, as well as unknown proteins A, B and C but not D. (**B**) CCRF-CEM cells inducibly express BirA*-HA alone or as a fusion with GFI1. Biotin-treated CCRF-CEM cells transduced with empty vector (lanes 1-2), BirA*-HA (lanes 3-4), or GFI1-BirA*-HA (lanes 5-6) were treated with doxycycline (Doxy) (+) or vehicle control (-). Whole cell lysates (WCL) were resolved by SDS-PAGE and immunoblotted using HA antibodies. *α*-tubulin is shown as a loading control. (**C**) Biotinylation of GFI1-associated proteins LSD1 and CoREST is enriched using GFI1-BirA*-HA expressing CCRF-CEM cells relative to empty vector or BirA*-HA controls. Whole cell lysates prepared as in (B) were used for immunoblotting or were first precipitated using streptavidin Sepharose beads (SAv-P), and subsequently analyzed by immunoblotting with anti-HA, anti-LSD1 or anti-CoREST antibodies, or streptavidin-HRP (SAv:HRP) to detect biotinylation.

To identify the complete cohort of GFI1 proximity partners, biotinylated proteins were purified on streptavidin (SAv)-Sepharose beads and subjected to unbiased, proteome-wide LC-MS/MS (Figure 1A). Three replicates were performed for each condition (Vector, BirA* and GFI1-BirA*). A total of 502 interacting proteins were identified that demonstrated both increased mass spectrometry intensity in the presence of doxycycline compared to empty vector control and a BirA* vs GFI1-BirA* *P-*value of <0.05 (Supplemental Table S1). Proximity partners were analyzed as previously described (McClellan, Casey et al., 2019). A volcano plot showing fold change in average sum read intensities (log_2_ (GFI1-WT/BirA*)) relative to *P*-value (-log_10_ *P*-value) is shown in Figure 2A. Being covalently tethered to BirA*, the placement of GFI1 in the top right corner of the plot is an important quality control, signifying the most statistically significant *P*-value and most abundantly enriched protein in the data set. Among biotinylated proteins, known GFI1 partners LSD1 (KDM1A), CoREST (RCOR1), STAG1, BCL11A and HMG20B were enriched when comparing GFI1-BirA* to BirA* only (Figure 2A, shown in red). Among newly-identified GFI1 proximity partners, the strongest overall by fold-enrichment was for IKZF1/IKAROS (Figure 2A). To identify possible functional protein associations, we clustered the top 40 GFI1-proximate proteins using the STRING functional protein—protein association network (Szklarczyk, Gable et al., 2019). The majority of these proteins were annotated as being nuclear and involved in regulating gene expression (Figure 2B, red and blue circles respectively). Notably, STRING output linked GFI1 directly to LSD1 and CoREST, as expected (Figure 2B), but placed IKAROS more remotely, arguing against a direct interaction with GFI1 (Figure 2B).

**Figure 2.**
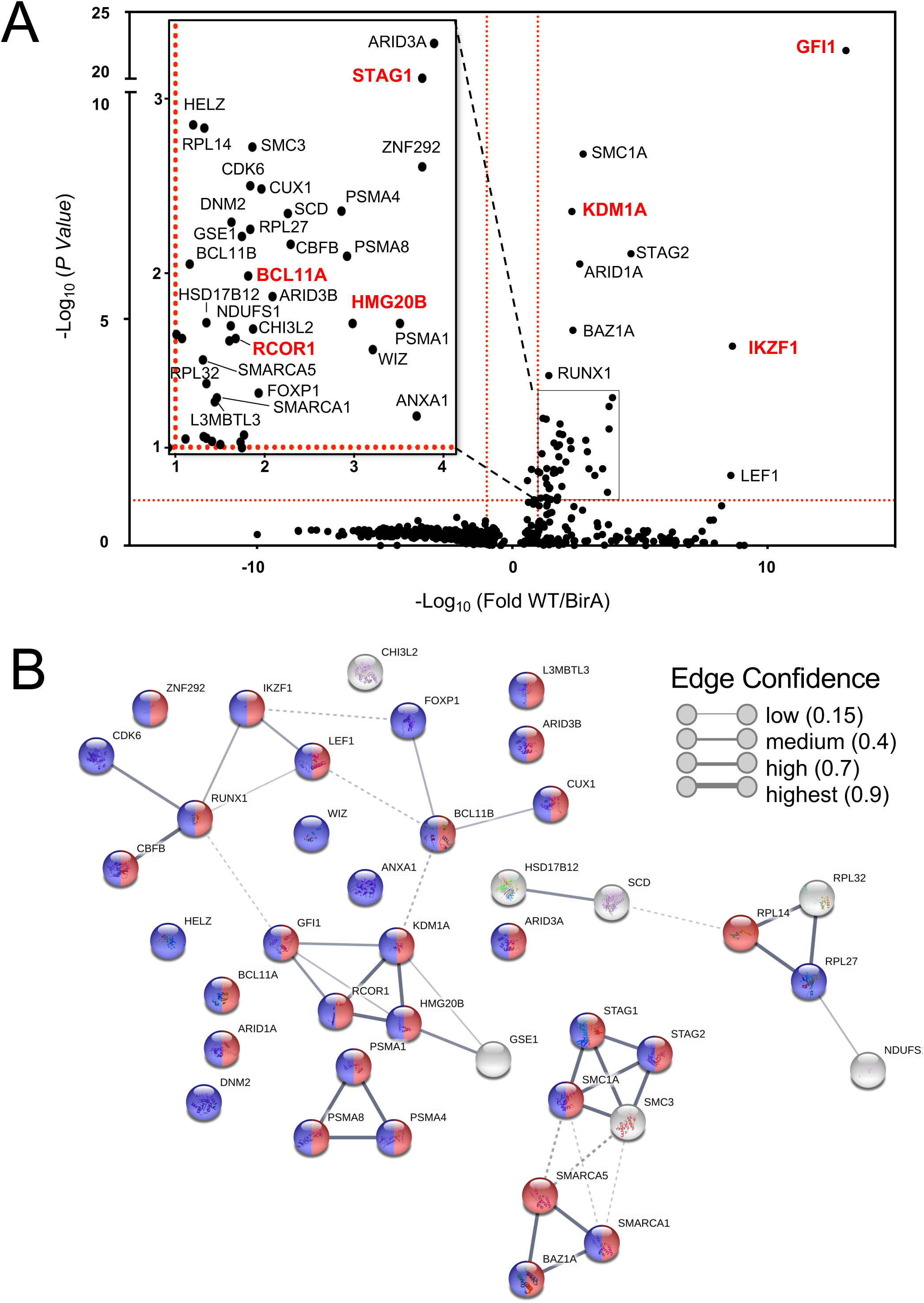
Top hits of GFI1 proximal partners identified by proteome-wide BioID labeling. (**A**) Volcano plot of proteins identified by BioID proximity labeling. Thresholds (red dashed lines) were fold-difference >2 and *P*-value <0.05. (**Inset**) Magnified portion of (A) depicting significant hits. Known GFI1 partners and GFI1 itself are labeled in red. (**B**) Genes labeled in (A) were subjected to STRING (version 10.5) functional protein association network analysis. Edges represent protein—protein associations. Blue indicates annotated nuclear proteins. Red indicates annotated proteins associated with gene regulation. The network was clustered using the MCL algorithm with inflation parameter 3. Dotted line indicates the proteins belong to different clusters.

### The GFI1—IKAROS interaction requires intact GFI1 and IKAROS DNA binding

We validated the proximity relationship between IKAROS and GFI1 through Western blotting. First, we tested if N-terminal GFI1 mutations known to block binding to LSD1 also block the ability to transfer biotin moieties to IKAROS. GFI1 mutations that block LSD1 binding did not affect IKAROS proximity labeling (Figure 3A). Additionally, GFI1 and IKAROS failed to interact using a variety of traditional co-immunoprecipitation conditions (Figure 3B and data not shown). One possible explanation for the discrepancy could be that GFI1 and IKAROS co-occupy nearby DNA binding sites at common target genes such that their proximity enables biotin moieties to be frequently transferred from GFI1-BirA* to IKAROS even though the two proteins do not interact directly in solution. Similarly, GFI1 and IKAROS could occupy sites distant from one another but brought together through interactions involving a shared protein complex. Both models predict that the proximity relationship between GFI1 and IKAROS would require both proteins to bind DNA and that a defect in DNA binding by either protein would attenuate the proximity relationship between them. Previous work on GFI1 and IKAROS has identified specific domains and mutants that control DNA binding activity (Kuehn, Boisson et al., 2016, Zarebski, Velu et al., 2008). We established these DNA-binding deficient mutations in GFI1 (N383S in rat GFI1, which corresponds to N382S in human GFI1) and IKAROS (N159A) and used them to interrogate the GFI1—IKAROS proximity relationship. GFI1-N383S-BirA* reduced proximity labeling of wild type IKAROS (Figure 3C). Likewise, we observed a comparable reduction in proximity labeling of IKAROS-N159A when when tested with wild type GFI1-BirA*. When both DNA-binding deficient mutants were combined, the proximity relationship between GFI1 and IKAROS is abolished (Figure 3C). These results indicate that the proximity relationship between GFI1 and IKAROS relies upon their shared ability to bind DNA and suggests a mechanism for GFI1 and IKAROS to cooperate to control a common set of genes through near or distant regulatory regions.

**Figure 3.**
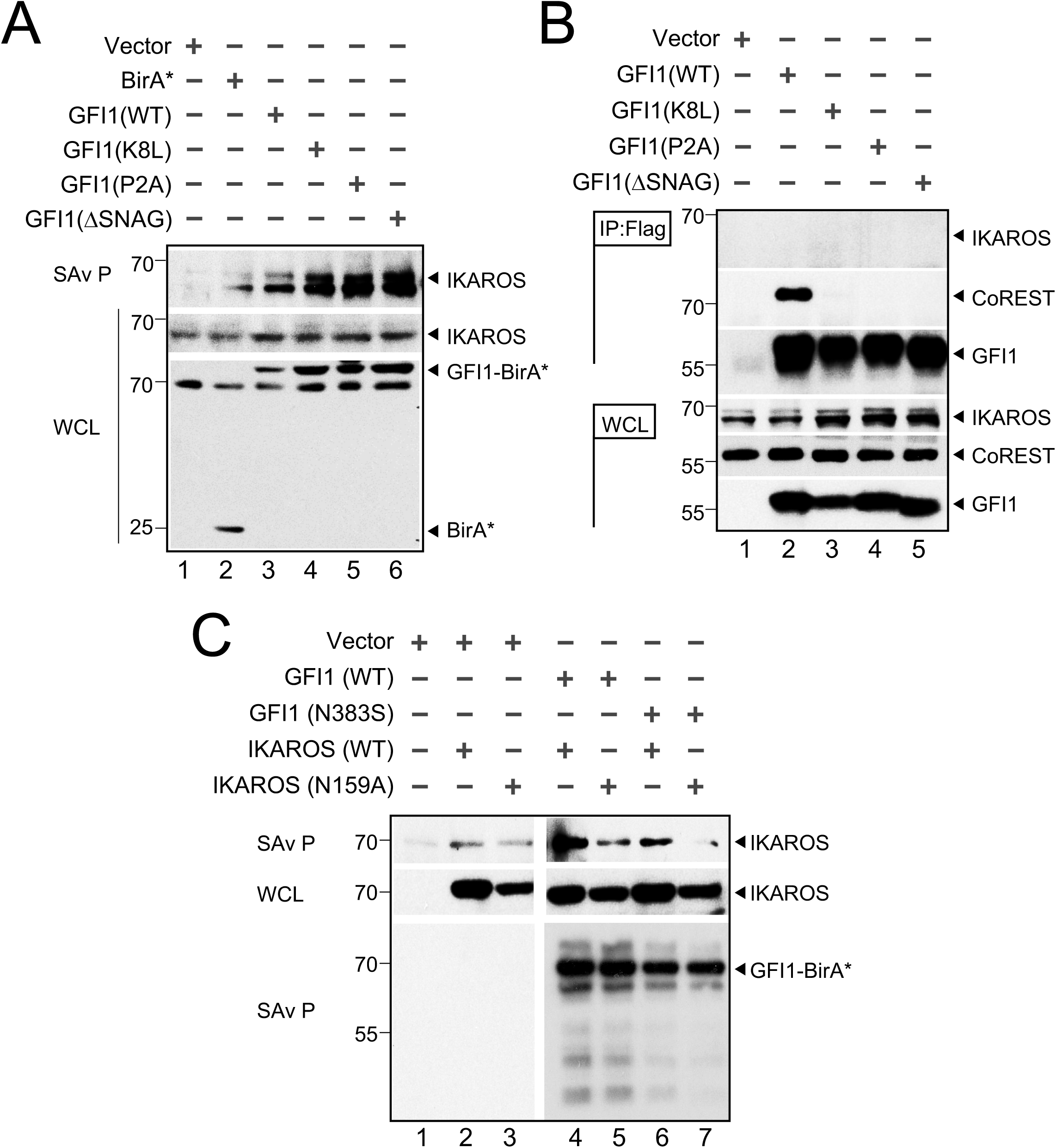
The GFI1—IKAROS interaction requires DNA binding. (**A**) Validation and non-LSD1 binding dependence of the GFI—IKAROS proximity relationship. Biotinylated proteins in whole cell lysates from CCRF-CEM cells transduced with the indicated constructs and treated as in Figure 1B were isolated with streptavidin beads and immunoblotted with anti-IKAROS and anti-HA antibodies targeting the epitope tag in GFI1-BirA* fusion protein variants. (**B**) GFI1—IKAROS binding is not observed in traditional coprecipitation methods. Lysates from CCRF-CEM cells inducibly expressing wild type GFI1-3*×*FLAG or variants (K8L,P2A and *Δ*SNAG) were immunoprecipitated with anti-FLAG antibody and Protein G-Sepharose. The presence or absence of IKAROS, CoREST, or GFI1 in the precipitate was detected by immunoblotting with anti-IKAROS, anti-CoREST or anti-FLAG antibodies. 2% input whole cell lysate (WCL) is shown as a control. (**C**) The proximity relationship between GFI1 and IKAROS requires DNA binding activity of both proteins. HEK293T cells were transiently transfected with empty vector or GFI1-BirA*-HA (wild type or N383S) together with human IKAROS-3*×*FLAG (wild type or N159A) expression constructs. IKAROS and control GFI1 biotinylation was monitored by precipitation with streptavidin beads and IKAROS immunoblotting as described in panel A. A GFI1 immunoblot is shown to confirm equivalent precipitation with streptavidin-Sepharose beads.

#### GFI1 and IKAROS associate with common genes, including genes associated with T cell development

To further study the interplay between GFI1 and IKAROS, we conducted ChIP-Seq using CCRF-CEM cells expressing 3*×*FLAG-tagged GFI1 or IKAROS (GFI1-3*×*FLAG or IKAROS-3*×*FLAG) under doxycycline-inducible control. FLAG immunoblotting confirmed inducible and comparable expression of the two proteins (Figure 4A). Two replicates each for GFI1 and IKAROS ChIP were performed. Sequencing of the ChIP material resulted in 25,674 total GFI1 and 52,759 IKAROS peaks. The signals from input, GFI1 ChIP-Seq and IKAROS ChIP-Seq replicates were highly correlated, with Spearman correlation R-values >0.8. Further, the GFI1 and IKAROS ChIP-Seq signals were also highly correlated, with R values >0.75. (Supplementary Figure S1A). Principal component analysis of the bound peaks also indicated that the inputs and ChIP replicates from the two cells lines were more similar to each other compared to the other samples (Supplementary Figure S1B). Approximately 80% of GFI1-bound peaks overlap (at least one bp) with peaks bound by IKAROS (Figure 4B), suggesting that they regulate common targets. To investigate this more closely, we centered binding peaks from either or both ChIP experiments (57,841 peaks) by peak summit, and arranged them as a heatmap from strongest to weakest GFI1 binding (Figure 4C, left side) and IKAROS binding (Figure 4C, right side), indicating an overall correlation between the strength of GFI1 and IKAROS binding. Motif analysis using all identified peaks showed that GFI1 and IKAROS shared four of the top five most significant binding motifs (Supplementary Figure S1C), again suggesting common gene binding. Among the top shared motifs is FLI1, an Ets transcription factor (consensus TTCC or GGAA reverse complement). Out of the top 20 enriched motifs for the two datasets, 14 and 15 Ets transcription motifs were present (Supplementary Figure S1C and data not shown). IKAROS is known to interact with Ets motifs (Zhang, Jackson et al., 2011).

**Figure 4.**
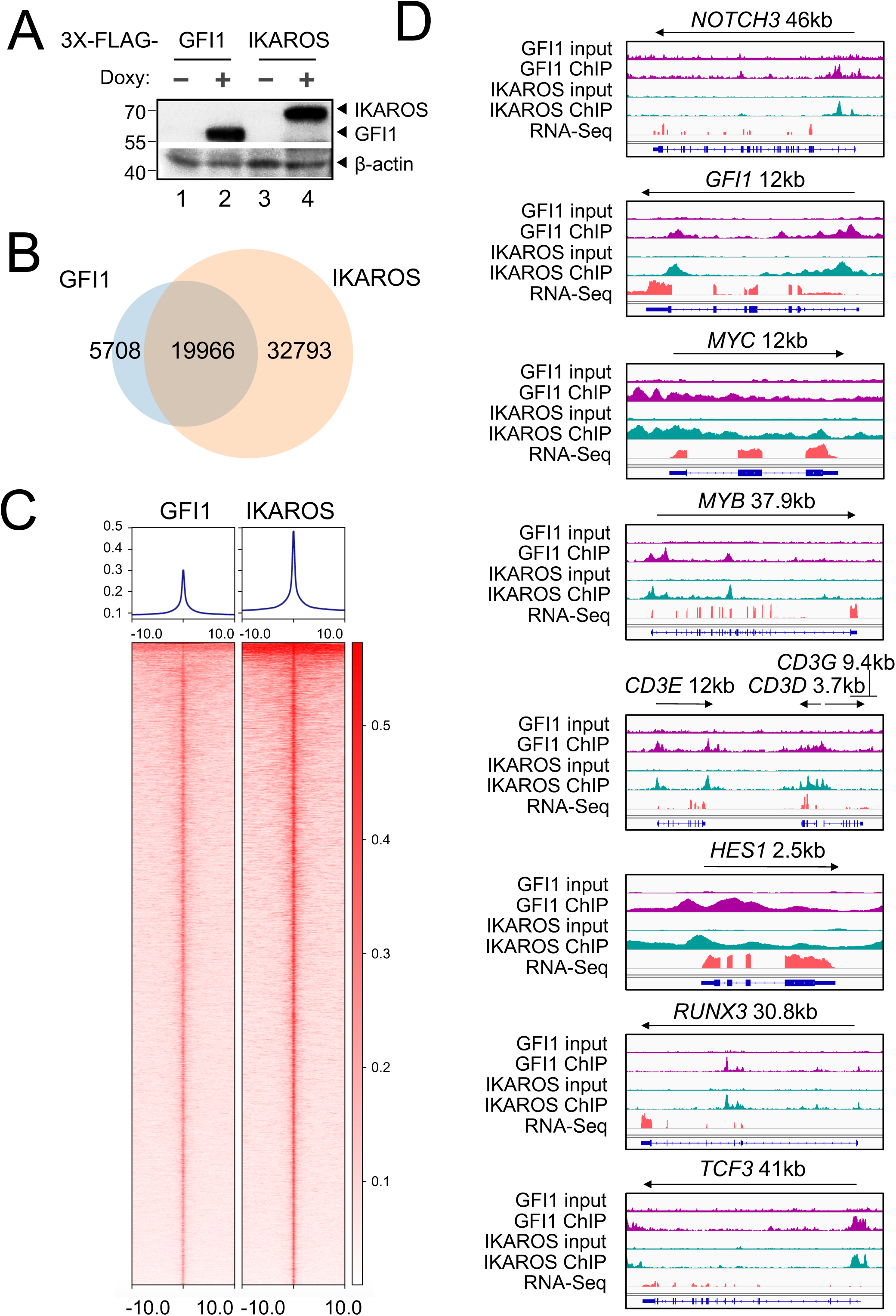
Identification of GFI1 and IKAROS targets in CCRF-CEM cells. (**A**) Validation of GFI1 and IKAROS expression in CCRF-CEM cell lines by Western blotting. CCRF-CEM cells expressing GFI1-3*×*FLAG and IKAROS-3*×*FLAG were used for ChIP-Seq. (**B**) Venn diagram showing enumeration of unique and shared GFI1 and IKAROS ChIP-Seq peaks. (**C**) Left: GFI1 ChIP-Seq heatmap showing all peak sites, centered by the ChIP peak summit and arranged from strongest to weakest. Right: the same set of peak summits in the same order shown as an IKAROS binding heatmap. (**D**) Example Integrative Genomics Viewer (IGV) tracks of GFI1 ChIP-Seq and input controls (purple), IKAROS ChIP-Seq and input controls (teal), together with previously published RNA-Seq (peach) of CCRF-CEM cells. *NOTCH3*, *GFI1*, *MYC*, *MYB*, *CD3*, *HES1*, *RUNX3* and *TCF3* tracks are shown.

GFI1 and IKAROS play crucial roles in hematopoiesis, including in T cells (Georgopoulos, Bigby et al., 1994, Shi, Kalupahana et al., 2013, Tinsley, Hong et al., 2013). Consistently, gene ontology (GO) analysis of genes within 6 kb of the GFI1- and IKAROS-bound peaks (using GREAT v4.0.4) identifies hematopoiesis and TCR recombination as common terms (Supplemental Table S2). Example bound targets are shown in Figure 4D, including genes encoding key transcription factors required for T cell development such as *GFI1* itself, *MYC*, *MYB*, *HES1, RUNX3* and *TCF3*. Other examples include *NOTCH3* and genes in the *CD3* cluster. Figure 4D also shows previously published RNA-Seq data (Quentmeier, Pommerenke et al., 2019) indicating that these genes are all expressed in CCRF-CEM cells.

### GFI1 and IKAROS positively regulate a subset of target genes

We used bulk RNA-Seq with GFI1-3*×*FLAG CCRF-CEM cells to identify changes in gene expression associated with either ectopic GFI1 expression or CRISPR-mediated IKAROS knockout. For knockout, we transfected GFI1-3*×*FLAG cells with CRISPR RNPs containing Cas9 protein, fluorescently-conjugated tracrRNA and either a nonspecific or *IKAROS*-specific sgRNA. Successfully transfected cells were sorted on the following day and cultured for an additional 9 days before treatment with doxycycline or vehicle for 24 hrs (see methods). This resulted in robust knockout using specific but not control RNPs (Figure 5A). A similar knockout strategy targeting GFI1 resulted in lethality in CCRF-CEM cells (not shown), underscoring its critical pro-survival role, and for this reason was not pursued further. Three independent replicates were performed for each of the four conditions, with the exception of vehicle-treated IKAROS knockout with two replicates. Between 21.7 and 28.2 million RNA-Seq reads were generated for each condition, wherein 88.6 to 90.3% of these aligned uniquely to the human *Hg38* reference genome. Approximately 99% of the reads within coding regions aligned to the correct strand (Supplemental Table S3). Hierarchical clustering of the top 500 most significantly differentially regulated genes across all the conditions (ranked based on *P*-value) revealed groups of genes repressed and activated by the different conditions. For example, a large number of genes are de-repressed with IKAROS knockout and/or repressed with GFI1 overexpression (Figure 5B, clusters 1, 2 ,3). Another group, not strongly affected by GFI1, was repressed with IKAROS knockout (cluster 5). However, a small group of 38 genes (cluster 4) were induced by GFI1 overexpression (Figure 5B). This same group of genes was also repressed with IKAROS knockout. Interestingly, when ectopic GFI1 expression and IKAROS knockout were combined, the induction of these genes was significantly blunted, suggesting IKAROS is required for GFI1-mediated transactivation (Figure 5B). These genes include *RAG1* and *NOTCH3* (Supplemental Table S4). Intersecting this set of positively regulated genes with the ChIP-Seq data reveals that 68.4% (26 of 38 total genes in cluster 4) have peaks for GFI1 and IKAROS located <10 kb from their transcription start sites, strongly suggesting they are direct targets of both GFI1 and IKAROS (Supplemental Table S4). 34.2% of these genes (13/38) show promoter binding (<500 bp from TSS) of both proteins. RNA-Seq genome tracks for *NOTCH3* are shown in Figure 5C alongside the GFI1 and IKAROS ChIP-Seq reads. Strong and overlapping GFI1 and IKAROS peaks are present in the 5’ region of the gene. In the RNA-seq, *NOTCH3* is induced by ectopic GFI1 expression and inhibited by IKAROS knockout, while the combination significantly blunts induction by GFI1. No such changes were observed for *NOTCH1* or *NOTCH2*, which show some GFI1 and IKAROS association but constitutive expression, or for *NOTCH4*, which shows no binding and is not expressed in CCRF-CEM cells (Supplementary Figure 2).

**Figure 5.**
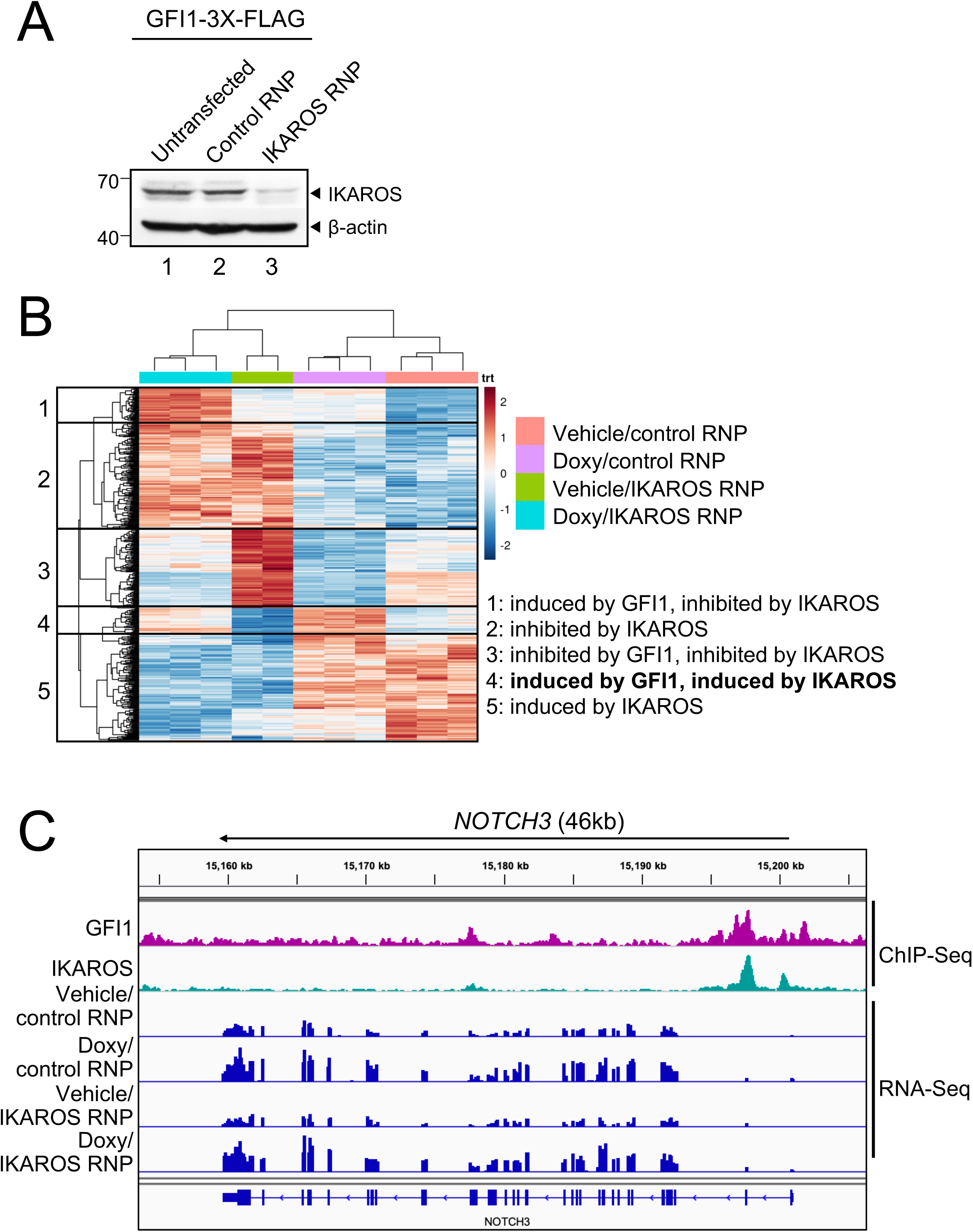
GFI1 and IKAROS positively regulate a subset of target genes. (**A**) IKAROS Western blot of untransfected CCRF-CEM-GFI1-3*×*FLAG cells, cells electroporated with ATTO550-conjugated control or IKAROS-specific RNPs. 24 hr post-transfection, ATTO550^+^ cells were sorted and cultured for 9 additional days prior to preparation of lysates. β-actin is shown as a loading control. (**B**) RNA-Seq analysis was performed using the same control RNP- or IKAROS-specific RNP-transfected cells shown in (A) after treatment with vehicle or doxycycline (Doxy) for 24 hr. The top 500 differentially expressed genes (based on *P*-value) were subjected to hierarchical clustering and are shown as a heatmap. Gene groupings with distinct expression patterns (1-5) are highlighted. (**C**) IGV tracks displaying GFI1 and IKAROS ChIP-Seq and RNA-Seq read distribution for *NOTCH3*.

### GFI1 and IKAROS cooperatively regulate NOTCH3

A model in which GFI1 and IKAROS interact on DNA to induce the expression of specific genes predicts that overexpression of either protein should induce target gene expression. Both GFI1 and IKAROS interact with the *NOTCH3* gene. In addition, NOTCH3 activating mutations can trigger T-ALL development (Bernasconi-Elias et al., 2016, Choi et al., 2017, Tottone et al., 2019). For these reasons, and because its surface expression can easily be measured using flow cytometry, we used *NOTCH3* as an example target gene. We treated CCRF-CEM cells that inducibly express GFI1-3*×*FLAG with doxycycline and followed surface NOTCH3 expression as a time course using flow cytometry. NOTCH3 was low but detectable in untreated cells (Figure 6A, 0h), with expression increasing during the 48 hour treatment course. Expression increased both as measured by the percentage of positive cells (not shown), and as measured by mean fluorescence intensity (MFI). CCRF-CEM cells inducibly expressing IKAROS-3*×*FLAG were similarly able to increase cell surface expression of NOTCH3 (Figure 6B). We then replaced the GFI1 lentiviral construct with a truncation mutant in which the SNAG domain was deleted (*Δ*SNAG) or with a SNAG domain point mutant that no longer interacts with LSD1 to mediate transcriptional repression (P2A) (Grimes, Chan et al., 1996, Velinder, Singer et al., 2017). Unexpectedly, these mutant forms of GFI1 no longer augment surface NOTCH3 expression (Figure 6C). These results suggest that GFI1 and IKAROS collaborate to transactivate the *NOTCH3* gene, and that GFI1 does so, at least in part, though residues that are also important to recruit LSD1. To more directly determine the role of LSD1 in GFI1-mediated NOTCH3 induction in CCRF-CEM cells, we utilized the non-competitive LSD1 inhibitor, SP-2509 (Fiskus, Sharma et al., 2017, Inui, Zhao et al., 2017). Surprisingly, NOTCH3 expression significantly increased in cells treated with SP-2509 compared to DMSO vehicle control even without doxycycline treatment (Figure 6D). Doxycycline-induced GFI1 expression cooperated with SP-2509 to further increase NOTCH3 expression at 24 but not 48 hr (Figure 6D). We also obtained similar results using SUP-T1, a different T-ALL cell line. Doxycycline-induced GFI1 expression or LSD1 inhibition with SP-2509 elevated surface levels of NOTCH3 (Figure 7A).

**Figure 6.**
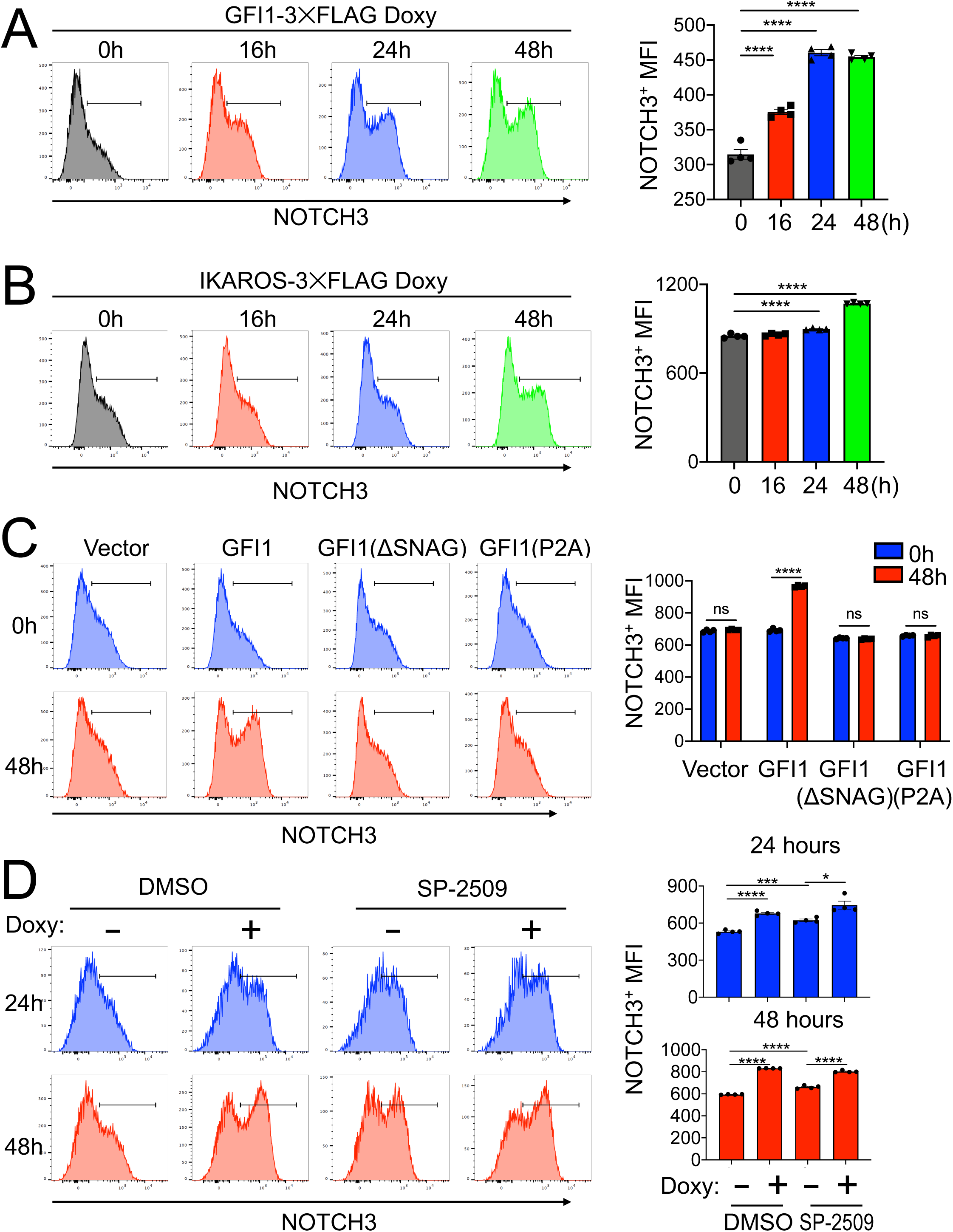
IKAROS cooperates with GFI1 to regulate cell surface NOTCH3 protein expression. (**A**) Left: example flow cytometry histograms showing NOTCH3 surface expression in CCRF-CEM-GFI1-3*×*FLAG cells treated with doxycycline (Doxy) for the indicated times. Untreated (0 hr) cells are shown as a control. Right: NOTCH3 MFIs from cells in the regions highlighted in the left panels were averaged from four experiments and plotted as a bar graph. Bar colors match the corresponding histograms. Error bars depict SEM. Ordinary one-way ANOVA was used for statistical analysis. In all bar graphs, values for individual data points are represented by closed shapes. (**B**) Left: CCRF-CEM-IKAROS-3*×*FLAG cells were treated with Doxy for the indicated times. Example NOTCH3 flow cytometry histograms are shown. Right: NOTCH3 MFIs from four independent experiments were averaged and plotted as in (A). Error bars depict SEM. Ordinary one-way ANOVA was used for statistical analysis. (**C**) CCRF-CEM cells transduced with empty vector (V), wild type GFI1-3*×*FLAG (GFI1), FLAG-tagged GFI1 lacking the N-terminal SNAG domain (*Δ*SNAG) or FLAG-tagged GFI1 with a SNAG domain point mutant no longer able to interact with LSD1 (P2A). Cells were treated with vehicle (blue) or Doxy (red) for 48 hr. Left: representative flow cytometry histograms for NOTCH3 cell surface expression. Right: average MFIs from four independent experiments. Error bars depict SEM. Two-way ANOVA was used for statistical analysis. (**D**) Flow cytometry of CCRF-CEM-GFI1-3*×*FLAG cells pre-treated with DMSO vehicle or LSD1 inhibitor (1 μM SP-2509, MedKoo Biosciences) for 20 hr and subsequently treated with Doxy for 24 or 48 hr. DMSO or SP-2509 was present continuously. Left: representative NOTCH3 flow cytometry. Right: MFIs within the NOTCH3-positive gates were averaged from four independent experiments and plotted as a bar graph. Error bars depict SEM. An unpaired T-test was used for statistical analysis.

**Figure 7.**
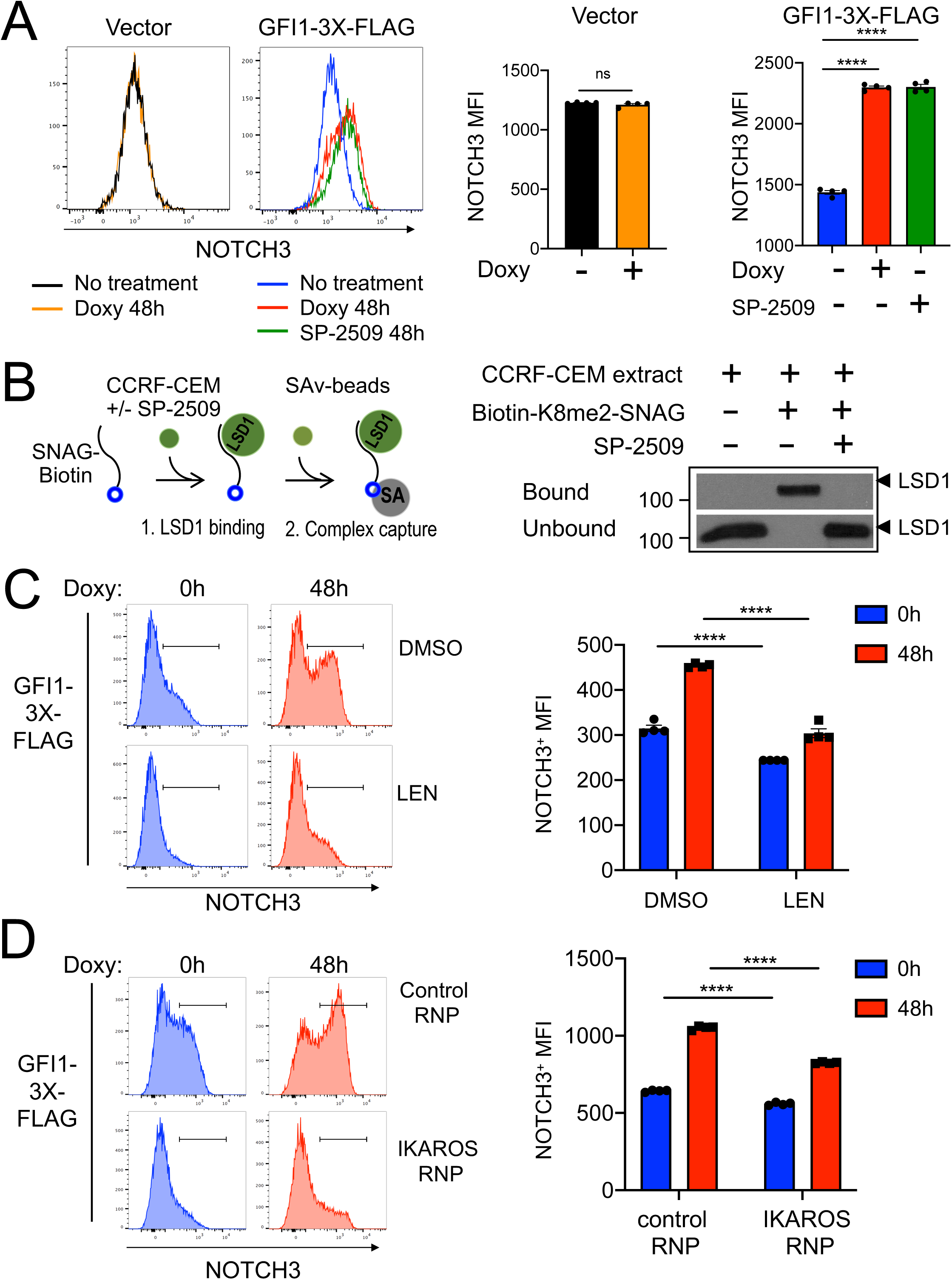
(**A**) Flow cytometry for NOTCH3 cell surface expression in SUP-T1cells transduced with vector or GFI1-3*×*FLAG inducible vectors and treated without or with doxycycline (Doxy), and SP-2509 for 48 hr. Left: representative NOTCH3 flow cytometry. Right: NOTCH3 MFIs were averaged from four independent experiments and plotted as a bar graph. Error bars depict SEM. An unpaired T-test was used for statistical analysis. In each bar graph, values for each independent test are represented by closed shapes. (**B**) SP-2509 abolishes the SNAG—LSD1 interaction. Biotinylated SNAG peptide demethylated at K8 was added to CCRF-CEM cell extracts with or without SP-2509 and incubated for 1hr prior to addition of streptavidin-Sepharose beads to collect biotinylated peptide and bound LSD1. LSD1 in the bound and unbound fractions were visualized by western blot using anti-LSD1 antibody. (**C**) Flow cytometry of CCRF-CEM-GFI1-3*×*FLAG cells pretreated with DMSO or 2 μM lenalidomide (LEN) for 16 hr before treatment with Doxy for 48 hr. DMSO or Lenalidomide was present continuously. Left: example NOTCH3 flow cytometry. Right: MFIs within the NOTCH3-positive regions shown at left were averaged from four independent experiments and plotted as a bar graph. Bar color corresponds to histogram shading on the left. Error bars depict SEM. Error bars depict SEM. Two-way ANOVA was used for statistical analysis. (**D**) Flow cytometry of CCRF-CEM-GFI1-3*×*FLAG cells electroporated with control RNP or IKAROS RNPs as in Figure 5A. Cells were treated with Doxy for 0 or 48 hr. Left: example NOTCH3 flow cytometry. Right: MFIs within the NOTCH3-positive regions shown on the left were averaged from four independent experiments and plotted as a bar graph. Error bars depict SEM. Two-way ANOVA was used for statistical analysis.

Notably, results obtained using SNAG domain mutants (*Δ*SNAG and P2A) are the reverse of those obtained with SP-2509, where the SNAG domain remains intact. We hypothesized this positive impact of SP-2509 could represent targeted disruption of the interaction between LSD1 and the SNAG domain to enable alternative interactions involving candidate co-activators. Small molecule inhibitors of LSD1 can disrupt binding between GFI1 and LSD1(Maiques-Diaz, Spencer et al., 2018). Further, we have previously shown the SNAG domain is sufficient for LSD1 binding and that dimethyl modification at lysine 8 (K8me2) of the SNAG domain strongly favors LSD1binding in an *in vitro* binding assay. We deployed this assay (Figure 7B, left panel) to test the impact of SP-2509 on SNAG—LSD1 binding. We find that SP-2509 completely abolishes binding between LSD1 and a biotinylated K8me2-SNAG peptide (Figure 7B, right panel). These results suggest the SNAG—LSD1 interaction can be modulated to enable competition between LSD1 and one or more unknown proteins for SNAG domain binding and regulation of GFI1-mediated transcriptional output.

We then tested the effect of IKAROS loss on GFI1-induced NOTCH3 expression using GFI1-3*×*FLAG CCRF-CEM cells. Lenalidomide is well-documented to rapidly and efficiently trigger degradation of IKAROS (Lu, Middleton et al., 2014). Induction of cell surface NOTCH3 expression was significantly impaired by Lenalidomide treatment compared with DMSO control (Figure 7C). Immunoblotting confirmed the rapid loss of IKAROS protein and unaltered expression of ectopic GFI1 in the presence of Lenalidomide (Supplementary Figure S3A-B). We obtained similar results using the same cells instead electroporated with IKAROS CRISPR RNPs (Figure 7D, Supp. Figure 3C-D). These results strongly suggest that IKAROS and GFI1 act in a cooperative fashion to transactivate a specific subset of genes which includes *NOTCH3*, and given the tumor promoting role of constitutively active NOTCH3 could provide a new direction for therapeutic development in T-ALL (Figure 8).

**Figure 8.**
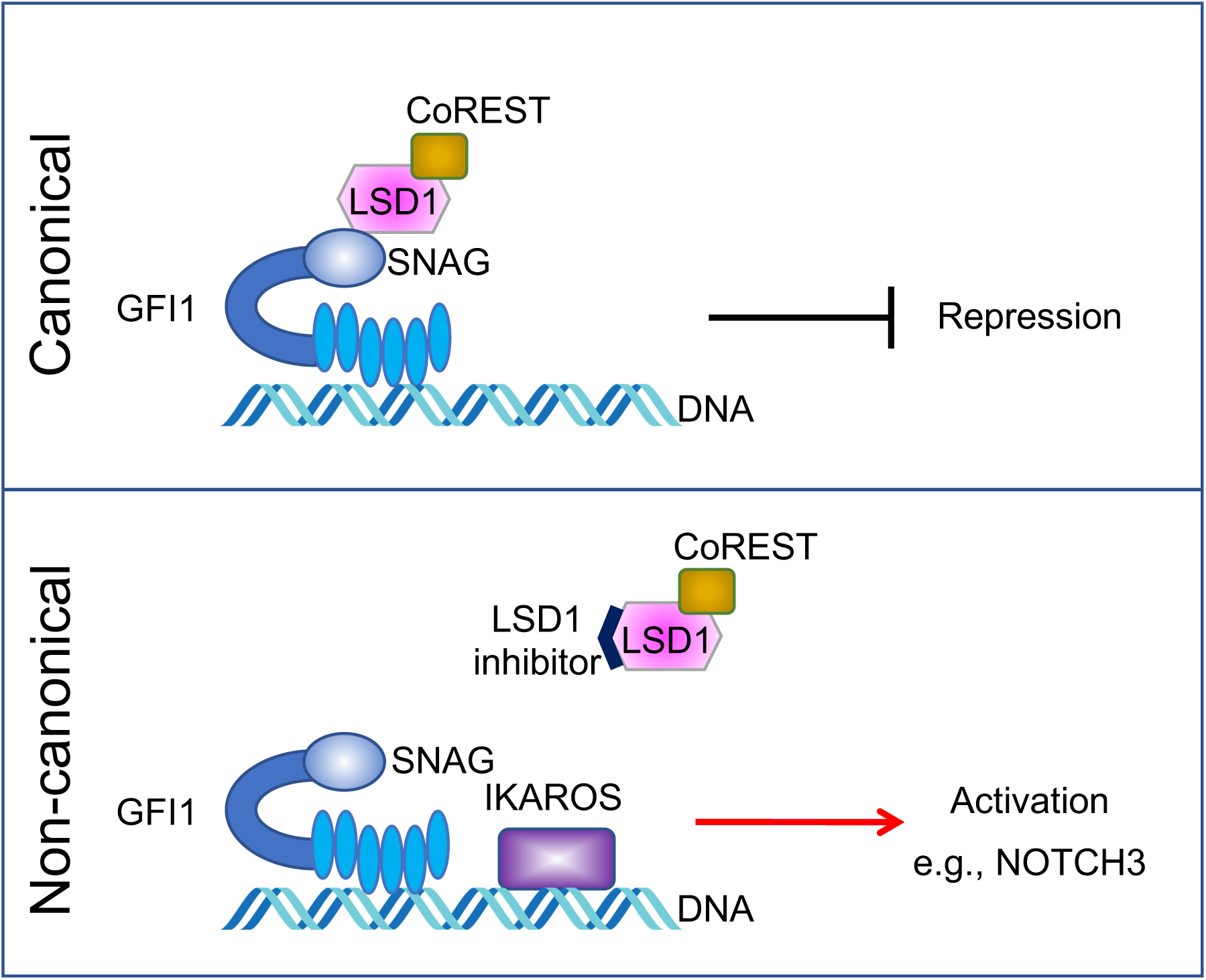
A proposed model of GFI1 mediated-noncanonical transcriptional activation. In the well-known canonical transcriptional regulation mechanism, GFI1 acts as a transcriptional repressor by recruiting LSD1/CoREST-containing complexes via its SNAG domain. In non-canonical transcriptional activation, GFI1 activates NOTCH3 and other hallmark T cell developmental genes with IKAROS. DNA binding by both proteins is required, as is an intact SNAG domain not bound by LSD1. An LSD1 inhibitor, which blocks SNAG—LSD1 binding enables transactivation of *NOTCH3* and similarly regulated genes via the GFI1—IKAROS partnership.

## Discussion

Identifying the molecular determinants of T cell development furthers understanding of T-ALL pathogenesis (Rothenberg & Taghon, 2005). The zinc finger transcription factor GFI1 plays critical roles in myeloid and lymphoid development and is an important pro-survival factor in T-ALL (Khandanpour et al., 2013, van der Meer, Jansen et al., 2010). The protein partnerships on which GFI1 depends for this role may offer potential therapeutic targets in T-ALL, but are as yet ill-defined. Here, using an unbiased quantitative proximity labeling approach, we identify GFI1 interacting proteins in T-ALL cells. Among these, IKAROS serves as a cooperating partner for GFI1-mediated gene regulation. Functional studies using T-ALL cells reveal a previously unrecognized role for GFI1 in transcriptional activation of genes that are among its targets, and several of which are involved in T cell development.

GFI1 and GFI1B have been largely described as a transcriptional repressors, and indeed can substitute for one another *in vivo*. Repression requires a highly conserved 20-amino acid N-terminal SNAG domain capable of recruiting LSD1 complexed with CoREST (Saleque, Kim et al., 2007). In previous studies, point mutations or truncations in the SNAG domain that disrupt GFI1/1B—LSD1 binding impair GFI1/1B functions in multiple assays, pinpointing LSD1 as a central cofactor in GFI1/1B-mediated transcriptional repression and the establishment of downstream phenotypes (Grimes et al., 1996, McClellan et al., 2019, Saleque et al., 2007, Velinder et al., 2017). However, it is not clear whether GFI1/1B can regulate gene expression through other mechanisms, especially in cooperation with other factors. We applied BioID proximity labeling to identify GFI1-interacting proteins where the promiscuous biotin ligase (BirA*) (Kim, Jensen et al., 2016, Roux, Kim et al., 2012) is fused to GFI1. Relative to co-immunoprecipitation, this method has the advantage that transient and indirect interactions can be efficiently captured. We utilized this method together with human CCRF-CEM T-ALL cells to systematically label proteins spatially close to GFI1, identifying GFI1 itself and some 500 direct or indirect interacting partners. Among these are previously identified GFI1-interacting proteins HMG20B, STAG1 and components of the BHC complex including LSD1, CoREST, and GSE1. We segregated interacting proteins into functional groups and analyzed their network relationships. Through this analysis we identified IKAROS, encoded by the *IKZF1* gene, as a top hit. IKAROS is also a zinc finger transcription factor that plays multiple, key roles in hematopoiesis, including in the development of B and T cells (Georgopoulos et al., 1994, Georgopoulos, Moore et al., 1992, Hahm, Ernst et al., 1994). As with GFI1, IKAROS has also been linked to mouse and human T-ALL. In T-ALL, IKAROS functions as a tumor suppressor, and its deletion is a poor prognostic indicator for affected patients (Marcais, Jeannet et al., 2010, Sun, Crotty et al., 1999, Winandy, Wu et al., 1995). This impact on prognosis takes on a new dimension when considered in the context of the GFI1—IKAROS partnership.

GFI1 and IKAROS do not co-precipitate in conventional co-IP assays, suggesting their interaction is indirect and transient in nature. Biotin transfer between the two proteins depends on DNA binding but does not depend on the GFI1 SNAG domain. Because GFI1 and IKAROS are both sequence-specific DNA binding proteins, we performed ChIP-Seq to identify potential common target genes. This effort identified >25,000 GFI1-bound and >50,000 IKAROS-bound peaks, and with considerable overlap. The large number of IKAROS peaks is consistent with previous reports for IKAROS binding in human K562 and myeloma cell lines (Barwick, Neri et al., 2019, Consortium, 2012) and may be attributable to its ability to bind DNA in multimeric configurations(McCarty, Kleiger et al., 2003, Molnar, Wu et al., 1996). Approximately 80% (19966/25674) of GFI1 bound peaks overlap with IKAROS peaks, supporting the idea that the two proteins frequently coregulate target gene expression. Co-occupied targets include *GFI1* itself, as well as *NOTCH3*, the *CD3* gene cluster (*CD3E*, *CD3D* and *CD3G*), *TCF3*, *MYC*, *RUNX3*, *MYB* and *HES1*, each with fundamental roles in T-cell development. Interplay between GFI1 and other transcription factors has been described. For example, GFI1 and HOXA9 compete for overlapping binding sites at a distinct class of targets in AML (Velu, Chaubey et al., 2014). Unlike HOXA9, our results suggest that IKAROS cooperates with GFI1.

RNA-Seq gene expression profiling using CCRF-CEM cells with ectopic GFI1 expression and/or with IKAROS knockout revealed a group of genes positively regulated by both proteins. Among these genes are *NOTCH3* and *RAG1*, while other NOTCH family members *NOTCH1*, *NOTCH2*, and *NOTCH4* were not affected by enforced expression of GFI1. This indicates *NOTCH3* is the specific NOTCH gene targeted for induction by the GFI1—IKAROS partnership. Moreover, genes making up this group are enriched for GFI1 and IKAROS occupancy, strongly suggesting they are direct targets and that the positive effects of the GFI1—IKAROS partnership on transcription are also direct in nature.

In CCRF-CEM cells *NOTCH1-*activating mutation and *FBXW7* loss-of-function drive constitutive NOTCH signaling (O’Neil, Grim et al., 2007). Similarly, SUP-T1 cells carry a t(7;9) translocation driving expression of a truncated NOTCH variant to produce constitutive, ligand-independent NOTCH activity (Reynolds, Smith et al., 1987). Using both cell lines, we find that GFI1 and IKAROS promote NOTCH3 cell surface expression. Moreover, mutations in the GFI1 SNAG domain that block the interaction with LSD1 also render GFI1 incapable of inducing NOTCH3. It is not clear whether LSD1 itself is important for this activation (e.g., through demethylating repressive protein modifications such as H3K9 methylation), or whether interactions between the SNAG domain and some other positively-acting cofactor become dominant in the case of GFI1—IKAROS coregulated genes. Interestingly, SP-2509 and SNAG mutants, both of which disrupt GFI1—LSD1 binding, yield qualitatively opposite results in NOTCH3 cell surface expression. Because SP-2509 leaves SNAG domain primary structure intact, our findings could indicate that LSD1 and an as yet unidentified co-activator compete for SNAG domain binding in an IKAROS-dependent manner. To address this possibility, we used lenalidomide to acutely abolish IKAROS expression. Lenalidomide is a thalidomide derivative that is active in multiple myeloma by enabling targeted degradation of IKAROS (Lu et al., 2014). We find that acute IKAROS depletion blunts GFI1-mediated transactivation and cell surface expression of NOTCH3, consistent with GFI1 and IKAROS providing a platform for SNAG-dependent recruitment of a transcriptional co-activating principle, leading to transactivation of a gene expression program represented by the *NOTCH3* response.

Cumulatively, the results support a model in which the presence of IKAROS allows for activation of target genes by GFI1. This model is consistent with the identification of SWI/SNF, a chromatin remodeling complex associated with positive regulation of gene expression (Hirschhorn, Brown et al., 1992, Imbalzano, Kwon et al., 1994, Kwon, Imbalzano et al., 1994, Peterson & Herskowitz, 1992), as a GFI1 proximity partner. An activating potential for both IKAROS and GFI1 has been described previously.

IKAROS binds and activates *Cd3d* through an upstream enhancer region (Georgopoulos et al., 1994, Georgopoulos et al., 1992). CD3 proteins form a central component of the T cell receptor signaling complex (Ngoenkam, Schamel et al., 2018). Positive regulation of gene expression by GFI1 was found in mouse granulocyte-monocyte precursor cells (GMPs) undergoing a binary monocyte/granulocyte fate decision. In GMPs it was shown that GFI1 associates with granulocyte-specific targets such as *Per3* and *Ets1*, activating their expression as part of a broader granulocyte fate-specifying program. Concurrently, GFI1 binding and repression of monocyte-specific genes suppresses the monocyte program (Olsson, Venkatasubramanian et al., 2016). It seems reasonable to consider that concurrent activation and repression of opposing gene expression programs, such as those exemplified by the GFI1—IKAROS—LSD1 relationship could direct alternative outcomes in developmental hematopoiesis. More work is necessary to define the molecular mechanisms by which GFI1 switches between repressive and activating transcriptional potential to control these binary fate decisions.

## Materials and Methods

### Cells and Culture conditions

CCRF-CEM and SUP-T1 cells were purchased from ATCC. Both cell lines were cultured in RPMI 1640 medium, 10% fetal bovine serum, 2 mM GlutaMAX-I, 100 U/mL penicillin and 100 μg/mL streptomycin. All cell culture materials were purchased from ThermoFisher Scientific, Waltham, MA.

### Antibodies

Antibodies used for immunoblotting and immunoprecipitation were as follows: anti-LSD1 (CST, C69G12, 2184S); anti-CoREST (CST, D6I2U,14567S); anti-IKAROS (ThermoFisher Scientific, PA5-23728); anti-*β*-actin (Santa Cruz, C4, sc-47778); anti-*α*-tubulin (Santa Cruz, B-7, sc-5286); anti-FLAG (Sigma, M2, F1804); anti-HA antibody (Roche, 12CA5, 11583816001). Streptavidin-HRP was purchased from GE Healthcare (RPN1231).

### Constructs and Cloning

In-frame GFI1 fusion proteins with BirA* were created by subcloning GFI1 from constructs in our previous study (Velinder et al., 2017) into the *Eco*RI and *Bam*HI sites of the MCS-BioID2-HA vector (a gift from Kyle Roux, Addgene #74224). Subsequently, GFI1-BirA*-HA PCR products were cloned into the *Not*I and *Eco*RI sites of the pLVX-Tight-Puro vector (Clontech Laboratories, Inc). We used the rat GFI1 ortholog for our experiments, which is >99% conserved at the amino acid level with human GFI1. IKZF1/IKAROS constructs were based on the human form. The N383S derivative of rat GFI1 is analogous to the N382S mutation in human GFI1.

### Lentivirus Packaging

To generate lentiviral particles, HEK293T cells were transfected with lentiviral packing vectors, viral packaging (psPAX2) and viral envelope (pMD2G) constructs at a 4:2:1 ratio with 1 mg/mL polyethylenimine (PEI, linear MD 25,000 Da, Sigma, Cat# 408727). The ratio of total transfected DNA to PEI was 1:3 (1 µg DNA:3 µg PEI). After 24 hr, 20 mL of fresh cell culture medium was added and the cells were incubated for an additional 24 hr. Then the culture medium was replaced with virus collection medium (culture medium with 20mM HEPES). Viral supernatants were collected after a further 8 and 24hr. The two supernatants were combined and passed through a 0.45μm filter. Virus was directly used for infection or stored at -80°C.

### BioID proximity purification and proteomic analysis (MassIVE File Identifier MSV000086405)

CCRF-CEM cells were transduced with Tet-On lentiviral vectors (Lenti-XTM Tet-On Advanced, Clontech Laboratories, Inc) and selected with 500 μg/mL G418 to generate CCRF-CEM-Tet-On cells. CCRF-CEM-Tet-On cells were transduced with empty vector, BirA*-HA or GFI1-BirA*-HA viruses (pLVX-Tight-Puro, pLVX-Tight-puro-BirA*-HA or pLVX-Tight-Puro-GFI1-BirA*-HA, respectively) and selected with 0.5

μg/mL puromycin to generate CCRF-CEM cells expressing doxycycline-inducible GFI1-BirA* fusion proteins or BirA*. Cells were treated with 1 μg/mL doxycycline for 48 hr and 20 μM biotin for 16 hr. Cells were lysed in buffer (50 mM Tris-HCl pH 7.5, 500 mM NaCl, 0.5 mM EDTA, 1% Triton X-100) and incubated with streptavidin-Sepharose High Performance beads (Sigma, GE17-5113-01) on ice for 16hr. After washing the beads five times with lysis buffer on ice, proteins were eluted with 2*×* Laemmli Sample Buffer (LSB, 65.8 mM Tris-HCl pH 6.8, 26.3% (w/v) glycerol, 2.1% SDS, 0.01% bromophenol blue) and boiled for 10 min. Proteins were resolved by SDS-PAGE. Whole lanes of SDS-PAGE gel were cut out and subjected to LTQ Orbitrap Velos Pro ion-trap mass spectrometry (ThermoFisher Scientific, Waltham, MA). Peptides were detected, isolated, and fragmented to produce a tandem mass spectrum of specific fragment ions for each peptide. Protein identity was determined by matching protein databases with the acquired fragmentation pattern by the software program, Sequest (ThermoFisher Scientific, Waltham, MA). All databases include a reversed version of all the sequences and the data was filtered to a 1-2% peptide false discovery rate.

### ChIP-Seq (Gene Expression Omnibus Series record GSE160183)

The CCRF-CEM-Tet-On cells described above were infected with GFI1-3*×*FLAG or IKAROS-3*×*FLAG viruses (packaged with pLVX-Tight-puro-GFI1-3*×*FLAG or pLVX-Tight-puro-IKAROS-3*×*FLAG plasmids). Cells were selected with 1μg/mL puromycin to produce CCRF-CEM stable cell lines inducibly expressing GFI1-3*×*FLAG and IKAROS-3*×*FLAG. These cells were treated with 1μg/mL doxycycline for 24 hr. 20 million CCRF-CEM cells were crosslinked using a final concentration of 1% formaldehyde in the medium for 10 min. Crosslinking was quenched with 0.125 M glycine for 2 min. Cells were washed with ice-cold PBS and Farnham lysis buffer (5 mM PIPES pH 8.0, 85 mM KCl, 0.5% NP-40, 1 mM PMSF and 10μg/mL Aprotinin pH 8.0). Cells were resuspended in 1 mL RIPA lysis buffer (1X PBS, 1% NP-40, 0.5% sodium deoxycholate, 0.1% SDS, 1 mM PMSF and 10 μg/mL Aprotinin pH 8.0). Cells were sonicated to shear the DNA to between 200 and 500 bp. Fifty µL of sonicated chromatin was saved. The leftover samples were precipitated with anti-FLAG antibody which was prebound to Protein G Dynabeads (Thermo Fisher). The 50 µL of saved chromatin and immunoprecipitated DNA were de-crosslinked in a 65°C water bath overnight. De-crosslinked samples were purified using a Zymo ChIP DNA clean & concentrator (Zymo Research). Five to 10 ng of precipitated DNA was used for library construction using NEBNext ChIP-Seq Library Prep Reagent Set. Sequencing libraries (25 pM) were chemically denatured and applied to an Illumina HiSeq v4 single read flow cell using an Illumina cBot. Hybridized molecules were clonally amplified and annealed to sequencing primers with reagents from an Illumina HiSeq SR Cluster Kit v4-cBot (GD-401-4001). Following transfer of the flow cell to an Illumina HiSeq 2500 instrument (HCSv2.2.38 and RTA v1.18.61), a 50-cycle single-read sequence run was performed using HiSeq SBS Kit v4 sequencing reagents (FC-401-4002).

### ChIP-Seq analysis

SAM alignments were generated from Illumina Fastq files aligned to human hg38 genome using Novocraft’s novoalign aligner (http://www.novocraft.com) with the following parameters: -o SAM –r Random. Peak calling was then performed using macs2 (https://github.com/taoliu/MACS, v2.1.1.20160309) with the following settings: -g 2.7e9 –call -summit –f BAMPE –nomodel –B –SPMR –extsize 200. Generated bedgraph files were then transformed to bw format using UCSC bedGraphToBigWig application (v4). Heatmap clustering of ChIP-SEQ was carried out using deepTools (v3). Matrix was generated with computeMatrix application using the following parameters: computeMatrix -S input_1.bw input_2.bw -R peaks.bed–outFileName out.matrix–referencePoint center -a 10000 -b 10000 -bs 100 –sortRegions descend. Correlation heatmap was generated using the deep Tools plot Heatmap application. PCA analysis was performed using the deepTools plotPCA application. The peaks.bed was generated by combining peaks from two ChIP-Seq experiments. plotHeatmap application with default settings was then used to plot heatmap. Peak regions were further used for motif finding analysis, which was carried out using the findMotifGenome.pl application (v4.8.3, HOMER, http://homer.ucsd.edu/homer/motif/) with default settings. This resulted in between 92.6M and 86.4M reads, wherein between 81.8M and 73.6M (88.3 and 85.2%) uniquely mapped to the *Hg38* genome build.

### IKAROS/IKZF1 knockout

Nonspecific or *IKZF1*-specific CRISPR/Cas9 RNPs were generated from tracrRNA-ATTO550 (IDT), crRNA (IDT) and Cas9 protein (QB3 MacroLab, UC Berkeley) using commercial guidelines (IDT) and transfected into GFI1-3*×*FLAG CCRF-CEM cells by electroporation using the Neon transfection system 10 μL kit (ThermoFisher Scientific). Electroporation parameters were 1500V, 10 ms pulse width, 3 pulses. Transfection efficiency was measured by flow cytometry to detect the tracrRNA-ATTO550 positive cells. Approximately 95% transfection efficiency was obtained. Cells were cultured for >9 days to confirm stable ablation of IKAROS protein by Western blotting. CRISPR guide RNAs were chosen targeting the 5’ exons of the *IKZF1* gene using the IDT predesigned CRISPR guide RNA database (https://www.idtdna.com/site/order/designtool/index/CRISPR_PREDESIGN): *IKAROS/IKZF1*: TCATCTGGAGTATCGCTTACagg; GACCTCTCCACCACCTCGGGagg; CTCCAAGAGTGACAGAGTCGtgg. CRISPR/Cas9 negative control crRNA (IDT, 1072544) was used as a control for transfection.

### RNA-Seq (Gene Expression Omnibus Series record GSE160183)

Total RNA was extracted from GFI1-3*×*FLAG CCRF-CEM cells treated with or without doxycycline for 24 hr, and with IKAROS or control CRISPR RNP knockout using the RNeasy Mini kit (Qiagen) and RNase-Free DNase Set (Qiagen). RNA integrity numbers (RIN) ranged from 9.4 to 9.9. Poly(A) RNA was purified from total RNA samples (100-500 ng) with oligo(dT) magnetic beads followed by library construction using the Illumina TruSeq Stranded mRNA Library Prep kit and TruSeq RNA UD Indexes. Purified libraries were qualified on an Agilent Technologies 2200 TapeStation using a D1000 ScreenTape assay. The molarity of adapter-modified molecules was defined by quantitative PCR using the Kapa Biosystems Kapa Library Quant Kit. Individual libraries were normalized to 1.30 nM in preparation for Illumina sequence analysis. NovaSeq 2*×*50 bp Sequencing_100 M Read-Pairs Sequencing libraries (1.3 nM) were chemically denatured and applied to an Illumina NovaSeq flow cell using the NovaSeq XP chemistry workflow. Following transfer of the flowcell to an Illumina NovaSeq instrument, a 2*×*51 cycle paired-end sequence run was performed using a NovaSeq S1 reagent Kit.

### RNA-seq analysis

Filtering and alignments were performed using the analysis pipeline developed by the Huntsman Cancer Institute (HCI) Bioinformatic Core Facility (https://huntsmancancerinstitute.github.io/hciRscripts/hciR_scripts.html). Briefly, fastq files were aligned using STAR aligned (v2.7.3a) with the following settings: --twopassMode Basic --outSAMtype BAM SortedByCoordinate --limitBAMsortRAM 64000000000 --outBAMsortingBinsN 100 --quantMode TranscriptomeSAM --outWigType bedGraph --outWigStrand Unstranded. The count matrix was then calculated using the Subread FeatureCounts function (v1.6.3) DESeq2 (v1.28.1) was used for differentially expressed genes analysis.

### *In vitro* SNAG—LSD1 binding assay

Biotinylated, lysine(K)-8 dimethylated SNAG peptide (Bio-K8me2-SNAG) was commercially synthesized and deployed as previously described in CCRF-CEM extracts in either the absence or presence of SP-2509 (Velinder et al., 2017). Biotinylated peptides were collected on streptavidin-Sepharose beads whose non-specific binding sites were blocked in 1% bovine serum albumin (BSA) in 1X phosphate buffered saline (PBS). LSD1 was detected by western blotting in pellets and supernatants.

## Data Availability

ChIP-Seq and RNA-Seq datasets are available at the Gene Expression Omnibus (GEO) website https://www.ncbi.nlm.nih.gov/geo/query/acc.cgi?acc=GSE160183. The following secure token has been created to allow reviewer access of the data while it remains in private status: gjircyuifxgxfaf. Should the manuscript by accepted for publication, the authors will direct GEO to release the data publicly. BioID mass spectrometry data were submitted to MassIVE with MassIVE File Identifier MSV000086405 at website: https://massive.ucsd.edu/ProteoSAFe/private-dataset.jsp?task=ff3728eabdb84663981aafe4e44df11b. The data is still in private status. The following reviewer login credentials will allow reviewers to access. Username :MSV000086405_reviewer; password: a.

## Acknowledgements

We thank F. Gounari and P. Ernst for critical reading of the manuscript. DNA sequencing and DNA oligonucleotides were synthesized by the University of Utah Health Sciences Center DNA/Peptide Synthesis Facility. We thank R. Tomaino at Taplin Mass Spectrometry Facility, Cell Biology Department, Harvard Medical School for assistance with LS-MS/MS and analysis. ChIP-Seq and RNA-Seq data reported in this study utilized the University of Utah Huntsman Cancer Institute High-Throughput Genomics and Bioinformatic Analysis Shared Resources. We thank C. Stubben at the Huntsman Cancer Institute Bioinformatics Shared Resource for assistance with submitting NGS sequencing data to GEO, and S.M. Osburn at the University Mass Spectrometry Facility for assistance with submitting mass spectrometry data to MassIVE.

## Author contributions

WS, DM, DB, AP and MC performed the experiments analyzed the data. JG analyzed ChIP-Seq and RNA-Seq data. MEE and DT conceived and supervised the study and interpreted data. BRC provided technical support. All authors were involved in the writing of the manuscript.

---

**Supplementary Figure S1.**
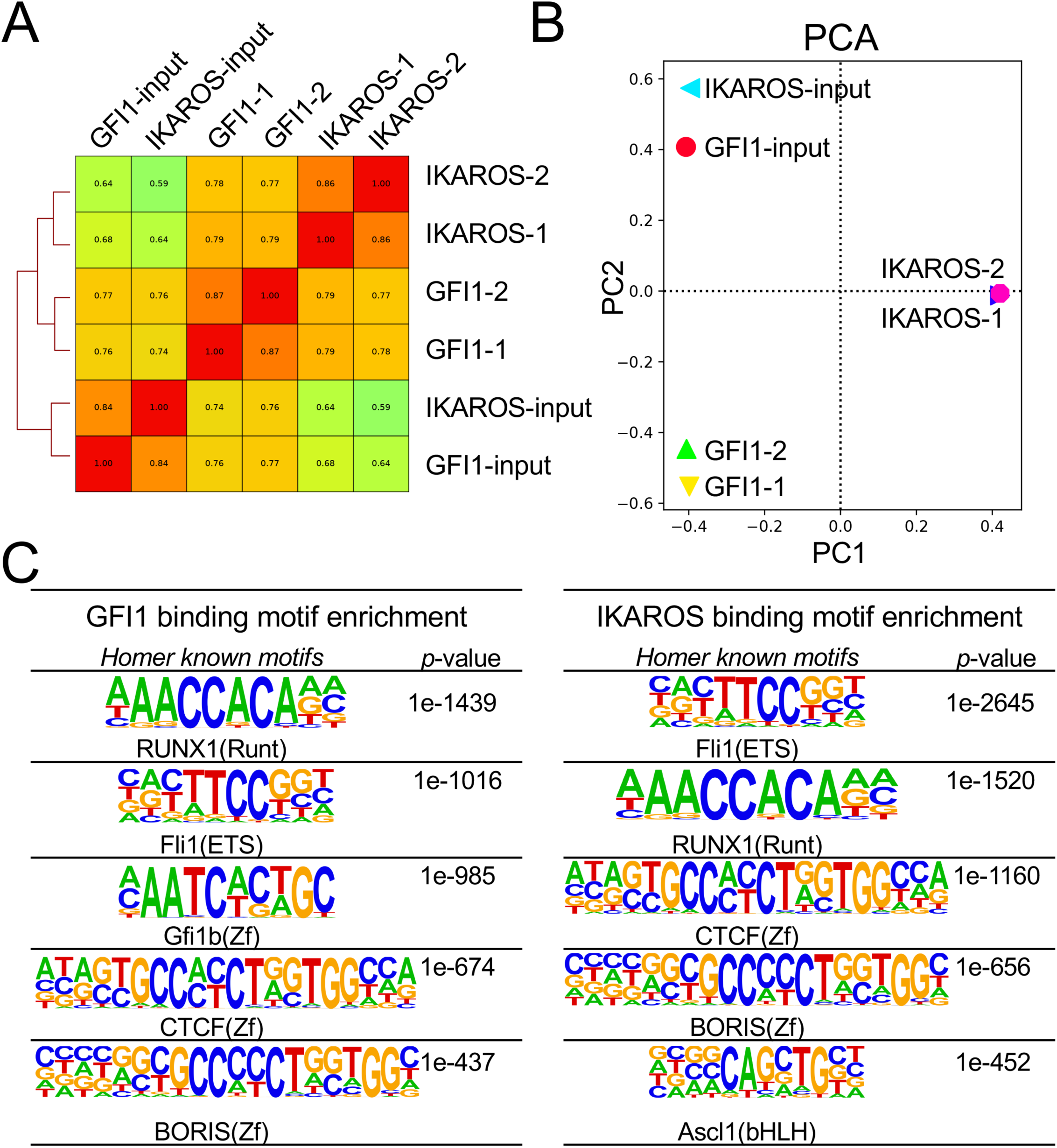
ChIP-Seq quality control data and motif analysis. (**A**) Correlation of GFI1 and IKAROS ChIP-Seq (and corresponding input) signaling. Spearman correlation R-values are displayed as a heatmap. (**B**) Principal component analysis (PCA) plot and cluster analysis of all ChIP-Seq samples. (**C**) Motif analysis (HOMER) showing enriched motifs within GFI1 and IKAROS ChIP-Seq peaks. The top 5 nonredundant motifs ranked by statistical significance are shown.

**Supplementary Figure S2.**
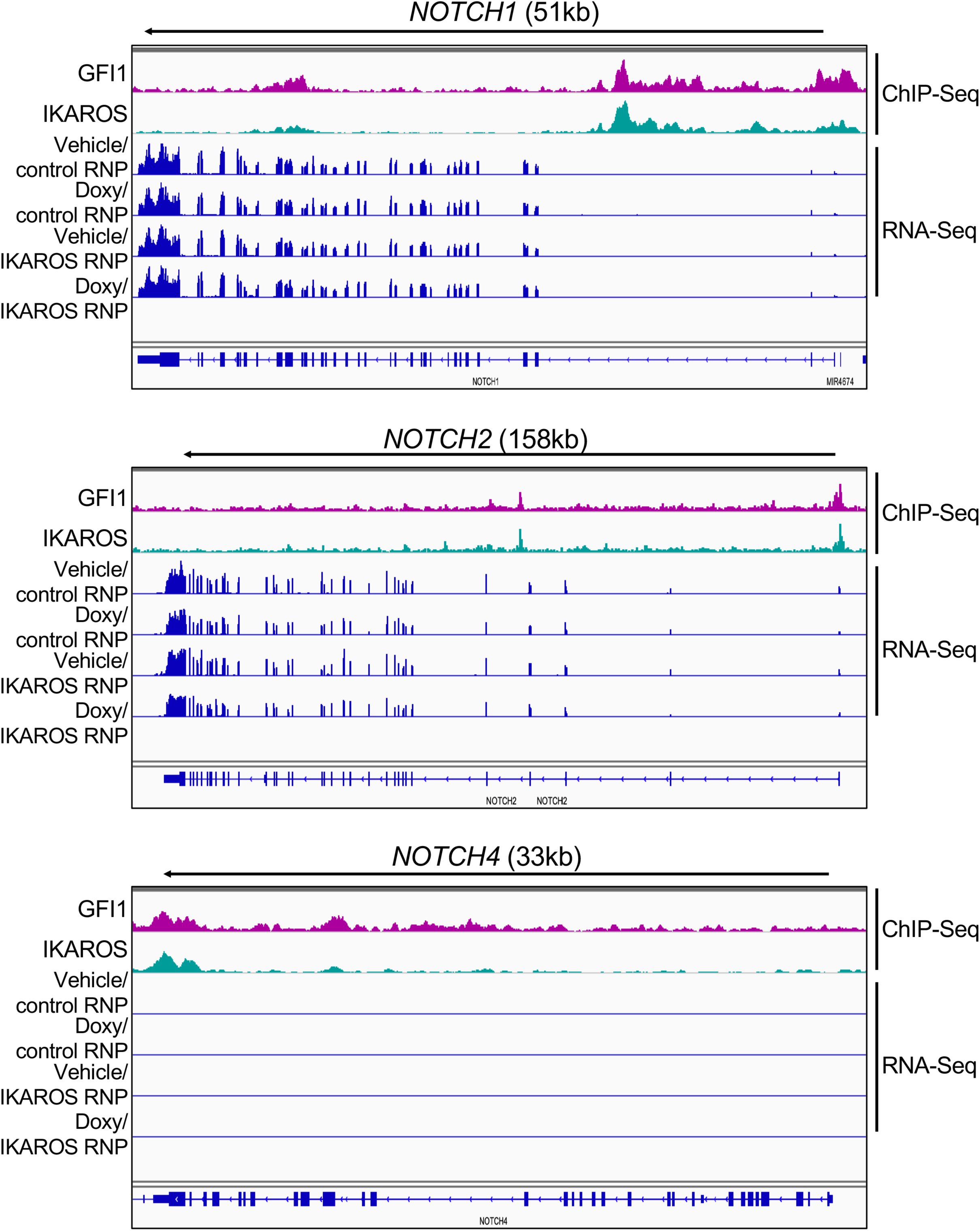
GFI1 and IKAROS binding and expression data for genes encoding other Notch family members. IGV tracks displaying GFI1 and IKAROS ChIP-Seq and RNA-Seq read distribution similar to Figure 5C, except for *NOTCH1*, *NOTCH2*, and *NOTCH4*.

**Supplementary Figure S3.**
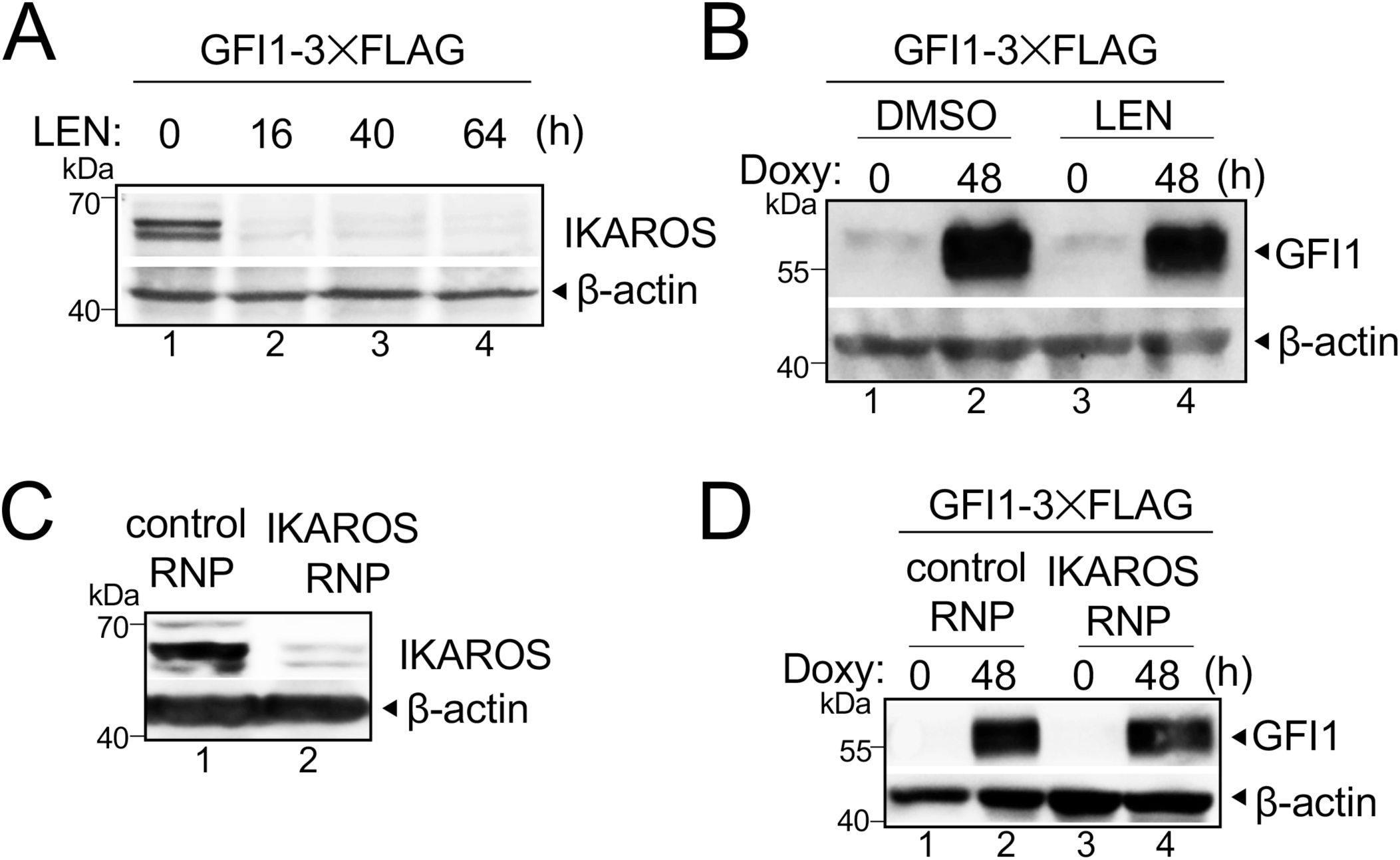
Lenalidomide treatment or CRIPSR RNP transfection rapidly depletes IKAROS protein while maintaining inducible GFI1 expression. (**A**) IKAROS Western blot of Lenalidomide (LEN)-treated CCRF-CEM GFI1-3*×*FLAG cells. Lysates were prepared from untreated (0 hr) cells, and over a 64 hr time course. β-actin is shown as a loading control. (**B**) Lysates from the same cells as in (A) were immunoblotted using GFI1 antibodies. (**C**) IKAROS Western blot using lysates from CCRF-CEM GFI1-3*×*FLAG cells electroporated with control or IKAROS RNP. Cells were prepared similarly as in Figure 5A. (**D**) Lysates from the same cells as in (C) were immunoblotted using GFI1 antibodies.

**Supplemental Table S1.** BioID mass spectrometry results (average intensity of WT<=VEC and three triplicates T-test (BirA* vs WT) *P*-value >/= 0.05, the data were deleted), 502 proteins.

**Supplemental Table S2.** Gene Ontology (GO) enrichment analysis of GFI1 and IKAROS ChIP-Seq peaks.

| Gene Ontology (GO) enrichment analysis with GF |  |  |
| --- | --- | --- |
| Rank | Term Name | P-Value |
| 1 | regulation of gene expression, epigenetic | 7.0764E-85 |
| 2 | regulation of hematopoietic progenitor cell differentiation | 3.4017E-68 |
| 3 | regulation of hematopoietic stem cell differentiation | 4.4564E-62 |
| 4 | negative regulation of gene expression, epigenetic | 4.3610E-52 |
|  | somatic diversification of immune receptors via germline |  |
| 5 | recombination within a single locus | 1.8269E-40 |
| 6 | V(D)J recombination | 2.1433E-39 |
| 7 | somatic diversification of immune receptors | 5.8618E-39 |
| 8 | regulation of megakaryocyte differentiation | 3.3875E-38 |
| 9 | DNA replication-dependent nucleosome assembly | 7.4252E-38 |
| 10 | positive regulation of viral life cycle | 2.0917E-33 |
| 11 | positive regulation of gene expression, epigenetic | 3.2093E-32 |
| 12 | chromatin silencing | 6.7381E-31 |
| 13 | positive regulation of myeloid leukocyte differentiation | 6.9143E-31 |
| 14 | beta-catenin-TCF complex assembly | 1.9700E-28 |
| 15 | chromatin silencing at rDNA | 3.6319E-28 |
|  | intrinsic apoptotic signaling pathway in response to DNA |  |
| 16 | damage | 4.8943E-27 |
|  | positive regulation of protein insertion into mitochondrial |  |
| 17 | membrane involved in apoptotic signaling pathway | 7.0662E-22 |

| Gene Ontology (GO) enrichment analysis with IKA |  |  |
| --- | --- | --- |
| Rank | Term Name | P-Value |
| 1 | regulation of hematopoietic progenitor cell differentiation | 1.1371E-61 |
| 2 | regulation of hematopoietic stem cell differentiation | 1.8733E-52 |
| 3 | B cell receptor signaling pathway | 4.4539E-40 |
|  | positive regulation of mitochondrial outer membrane |  |
| 4 | permeabilization involved in apoptotic signaling pathway | 7.3392E-34 |
| 5 | somatic diversification of immune receptors | 1.1375E-32 |
| 6 | regulation of T cell receptor signaling pathway | 3.9032E-26 |

\*The GO analysis was performed using GREAT tool (v 4.0.4) following its default settings.

s of GFI1 and IKAROS ChIP-SEQ peaks\*
| GFI1 binding peaks |  |  |
| --- | --- | --- |
| Fold Enrichment | Observed Gene Hits | Total Genes |
| 2.1208 | 209 | 248 |
| 2.7850 | 92 | 113 |
| 2.8136 | 86 | 103 |
| 2.5319 | 93 | 114 |
| 2.9966 | 31 | 32 |
| 3.6440 | 14 | 14 |
| 2.7361 | 36 | 38 |
| 2.2967 | 65 | 76 |
| 7.2147 | 29 | 32 |
| 2.0164 | 78 | 98 |
| 2.4171 | 70 | 80 |
| 2.4221 | 78 | 97 |
| 2.0172 | 45 | 50 |
| 2.1595 | 39 | 43 |
| 4.0207 | 35 | 36 |
| 2.1653 | 57 | 69 |
| 2.3287 | 27 | 29 |

| IKAROS binding peaks |  |  |
| --- | --- | --- |
| Fold Enrichment | Observed Gene Hits | Total Genes |
| 2.1139 | 107 | 113 |
| 2.0869 | 97 | 103 |
| 2.0451 | 35 | 35 |
| 2.079 | 37 | 37 |
| 2.0353 | 38 | 38 |
| 2.0342 | 36 | 36 |

**Supplemental Table S3.** RNA-Seq differential gene expression.

**Supplemental Table S4.**
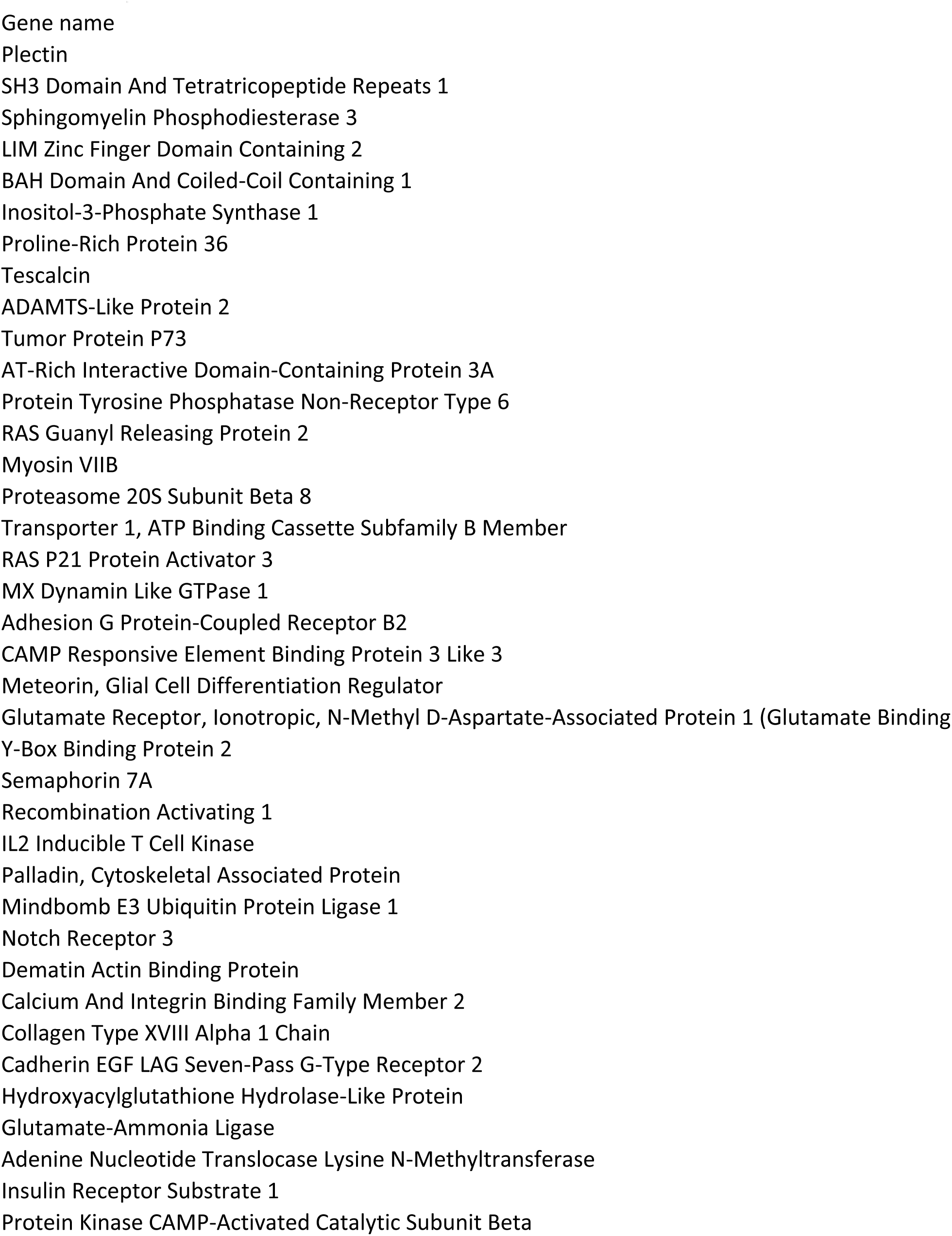
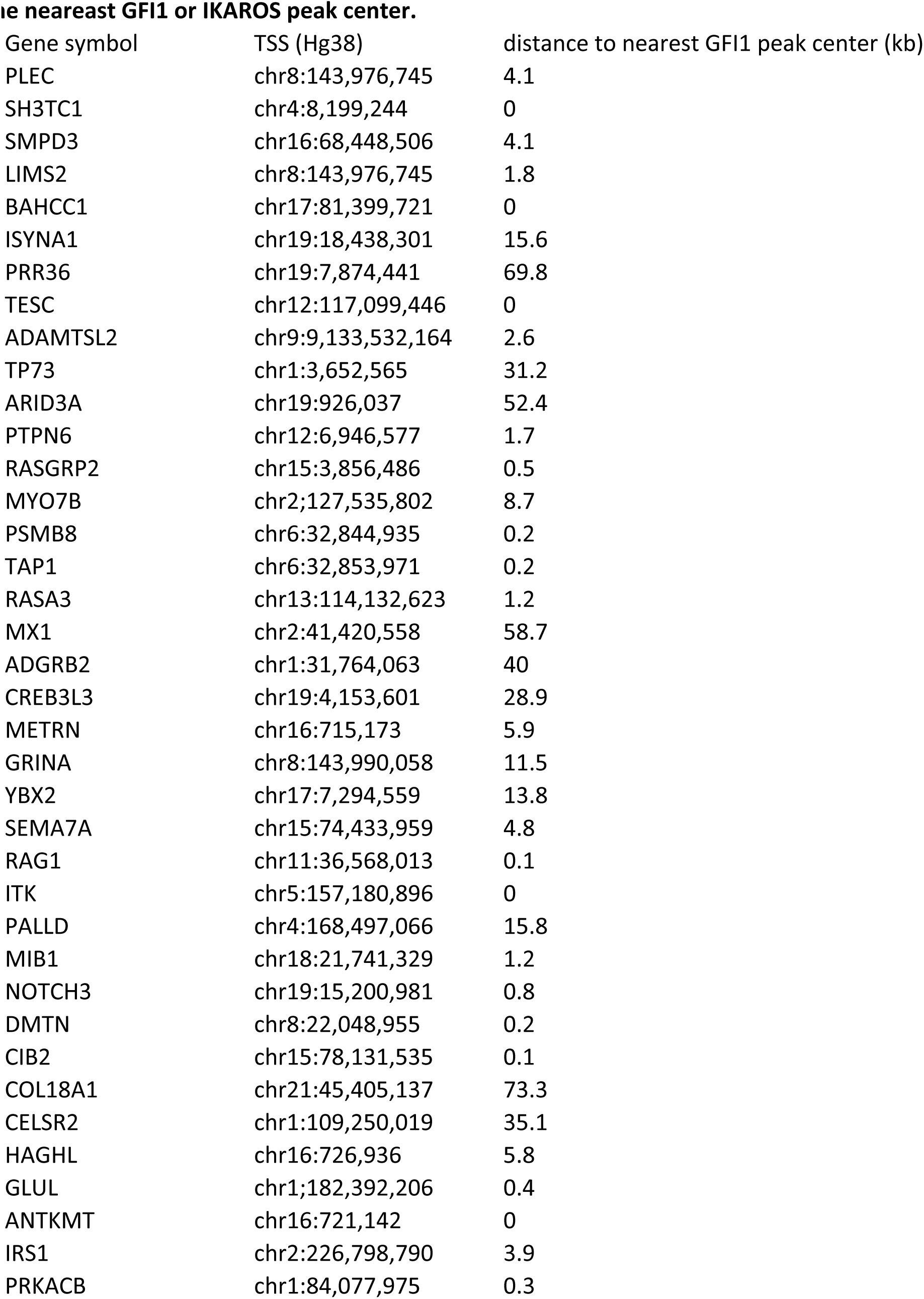
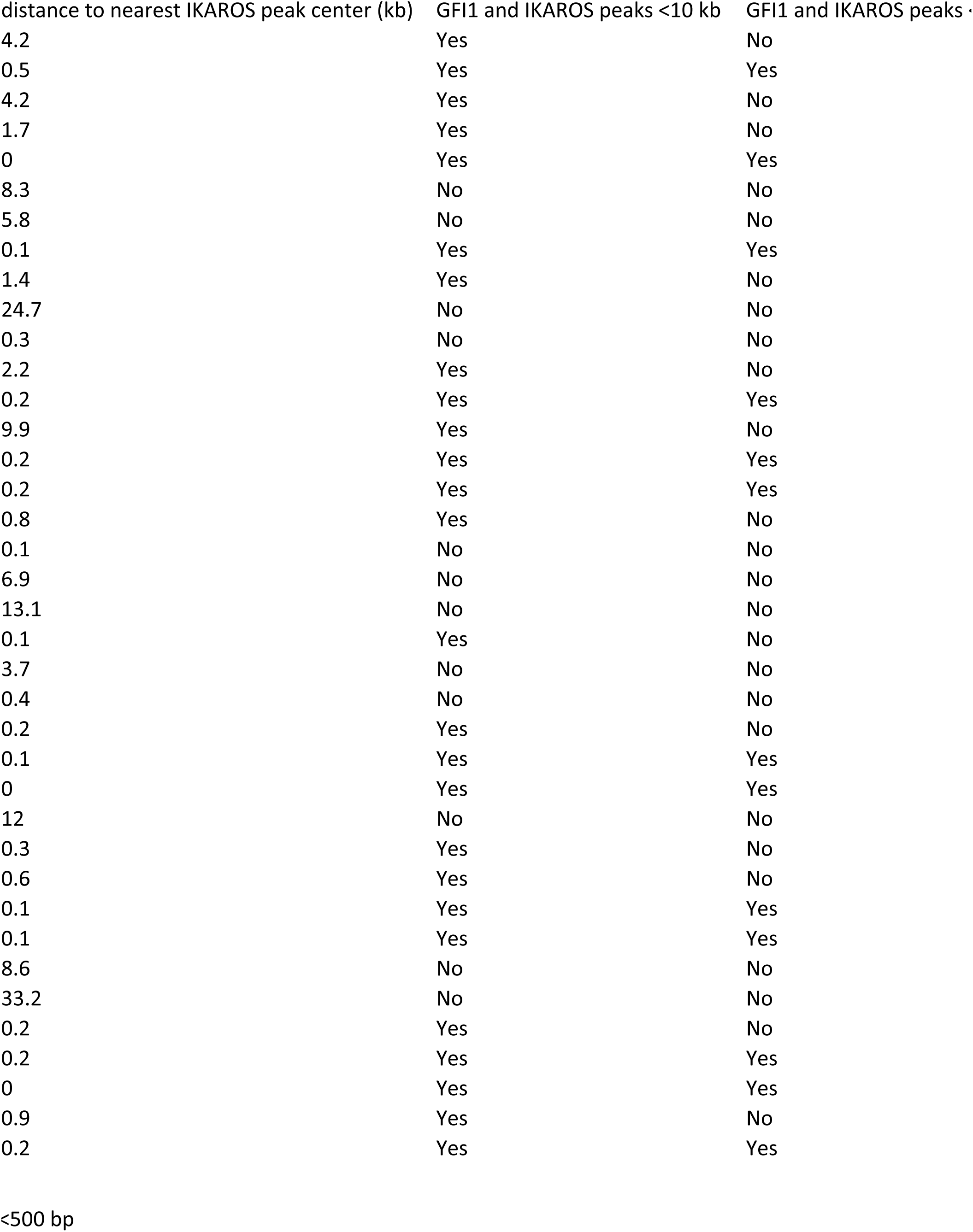
List of genes within cluster 4 of Figure 5B and distance from TSS to the nearest GFI1 or IKAROS peak center.

## References

Barwick BG, Neri P, Bahlis NJ, Nooka AK, Dhodapkar MV, Jaye DL, Hofmeister CC, Kaufman JL, Gupta VA, Auclair D, Keats JJ, Lonial S, Vertino PM, Boise LH (2019) Multiple myeloma immunoglobulin lambda translocations portend poor prognosis. Nat Commun 10: 1911

Bellavia D, Campese AF, Alesse E, Vacca A, Felli MP, Balestri A, Stoppacciaro A, Tiveron C, Tatangelo L, Giovarelli M, Gaetano C, Ruco L, Hoffman ES, Hayday AC, Lendahl U, Frati L, Gulino A, Screpanti I (2000) Constitutive activation of NF-kappaB and T-cell leukemia/lymphoma in Notch3 transgenic mice. EMBO J 19: 3337–48

Bellavia D, Campese AF, Checquolo S, Balestri A, Biondi A, Cazzaniga G, Lendahl U, Fehling HJ, Hayday AC, Frati L, von Boehmer H, Gulino A, Screpanti I (2002) Combined expression of pTalpha and Notch3 in T cell leukemia identifies the requirement of preTCR for leukemogenesis. Proc Natl Acad Sci U S A 99: 3788–93

Bernasconi-Elias P, Hu T, Jenkins D, Firestone B, Gans S, Kurth E, Capodieci P, Deplazes-Lauber J, Petropoulos K, Thiel P, Ponsel D, Hee Choi S, LeMotte P, London A, Goetcshkes M, Nolin E, Jones MD, Slocum K, Kluk MJ, Weinstock DM et al. (2016) Characterization of activating mutations of NOTCH3 in T-cell acute lymphoblastic leukemia and anti-leukemic activity of NOTCH3 inhibitory antibodies. Oncogene 35: 6077–6086

Choi SH, Severson E, Pear WS, Liu XS, Aster JC, Blacklow SC (2017) The common oncogenomic program of NOTCH1 and NOTCH3 signaling in T-cell acute lymphoblastic leukemia. PLoS One 12: e0185762

Consortium EP (2012) An integrated encyclopedia of DNA elements in the human genome. Nature 489: 57–74

Coustan-Smith E, Mullighan CG, Onciu M, Behm FG, Raimondi SC, Pei D, Cheng C, Su X, Rubnitz JE, Basso G, Biondi A, Pui CH, Downing JR, Campana D (2009) Early T-cell precursor leukaemia: a subtype of very high-risk acute lymphoblastic leukaemia. Lancet Oncol 10: 147–56

Ferrando A (2010) NOTCH mutations as prognostic markers in T-ALL. Leukemia 24: 2003–4

Fiskus W, Sharma S, Shah B, Portier BP, Devaraj SGT, Liu K, Iyer SP, Bearss D, Bhalla KN (2017) Highly effective combination of LSD1 (KDM1A) antagonist and pan-histone deacetylase inhibitor against human AML cells. Leukemia 31: 1658

Georgopoulos K, Bigby M, Wang JH, Molnar A, Wu P, Winandy S, Sharpe A (1994) The Ikaros gene is required for the development of all lymphoid lineages. Cell 79: 143–56

Georgopoulos K, Moore DD, Derfler B (1992) Ikaros, an early lymphoid-specific transcription factor and a putative mediator for T cell commitment. Science 258: 808–12

Golde TE, Koo EH, Felsenstein KM, Osborne BA, Miele L (2013) gamma-Secretase inhibitors and modulators. Biochim Biophys Acta 1828: 2898–907

Grimes HL, Chan TO, Zweidler-McKay PA, Tong B, Tsichlis PN (1996) The Gfi-1 proto-oncoprotein contains a novel transcriptional repressor domain, SNAG, and inhibits G1 arrest induced by interleukin-2 withdrawal. Mol Cell Biol 16: 6263–72

Hahm K, Ernst P, Lo K, Kim GS, Turck C, Smale ST (1994) The lymphoid transcription factor LyF-1 is encoded by specific, alternatively spliced mRNAs derived from the Ikaros gene. Mol Cell Biol 14: 7111–23

Hirschhorn JN, Brown SA, Clark CD, Winston F (1992) Evidence that SNF2/SWI2 and SNF5 activate transcription in yeast by altering chromatin structure. Genes Dev 6: 2288–98

Hock H, Hamblen MJ, Rooke HM, Schindler JW, Saleque S, Fujiwara Y, Orkin SH (2004) Gfi-1 restricts proliferation and preserves functional integrity of haematopoietic stem cells. Nature 431: 1002–7

Hock H, Hamblen MJ, Rooke HM, Traver D, Bronson RT, Cameron S, Orkin SH (2003) Intrinsic requirement for zinc finger transcription factor Gfi-1 in neutrophil differentiation. Immunity 18: 109–20

Hones JM, Botezatu L, Helness A, Vadnais C, Vassen L, Robert F, Hergenhan SM, Thivakaran A, Schutte J, Al-Matary YS, Lams RF, Fraszscak J, Makishima H, Radivoyevitch T, Przychodzen B, da Conceicao Castro SV, Gorgens A, Giebel B, Klein-Hitpass L, Lennartz K et al. (2016) GFI1 as a novel prognostic and therapeutic factor for AML/MDS. Leukemia 30: 1237–45

Imbalzano AN, Kwon H, Green MR, Kingston RE (1994) Facilitated binding of TATA-binding protein to nucleosomal DNA. Nature 370: 481–5

Inui K, Zhao Z, Yuan J, Jayaprakash S, Le LTM, Drakulic S, Sander B, Golas MM (2017) Stepwise assembly of functional C-terminal REST/NRSF transcriptional repressor complexes as a drug target. Protein Sci 26: 997–1011

Karsunky H, Zeng H, Schmidt T, Zevnik B, Kluge R, Schmid KW, Duhrsen U, Moroy T (2002) Inflammatory reactions and severe neutropenia in mice lacking the transcriptional repressor Gfi1. Nat Genet 30: 295–300

Khandanpour C, Krongold J, Schutte J, Bouwman F, Vassen L, Gaudreau MC, Chen R, Calero-Nieto FJ, Diamanti E, Hannah R, Meyer SE, Grimes HL, van der Reijden BA, Jansen JH, Patel CV, Peeters JK, Lowenberg B, Duhrsen U, Gottgens B, Moroy T (2012) The human GFI136N variant induces epigenetic changes at the Hoxa9 locus and accelerates K-RAS driven myeloproliferative disorder in mice. Blood 120: 4006–17

Khandanpour C, Phelan JD, Vassen L, Schutte J, Chen R, Horman SR, Gaudreau MC, Krongold J, Zhu J, Paul WE, Duhrsen U, Gottgens B, Grimes HL, Moroy T (2013) Growth factor independence 1 antagonizes a p53-induced DNA damage response pathway in lymphoblastic leukemia. Cancer Cell 23: 200–14

Kim DI, Jensen SC, Noble KA, Kc B, Roux KH, Motamedchaboki K, Roux KJ (2016) An improved smaller biotin ligase for BioID proximity labeling. Mol Biol Cell 27: 1188–96

Kuehn HS, Boisson B, Cunningham-Rundles C, Reichenbach J, Stray-Pedersen A, Gelfand EW, Maffucci P, Pierce KR, Abbott JK, Voelkerding KV, South ST, Augustine NH, Bush JS, Dolen WK, Wray BB, Itan Y, Cobat A, Sorte HS, Ganesan S, Prader S et al. (2016) Loss of B Cells in Patients with Heterozygous Mutations in IKAROS. N Engl J Med 374: 1032–1043

Kwon H, Imbalzano AN, Khavari PA, Kingston RE, Green MR (1994) Nucleosome disruption and enhancement of activator binding by a human SW1/SNF complex. Nature 370: 477–81

Lu G, Middleton RE, Sun H, Naniong M, Ott CJ, Mitsiades CS, Wong KK, Bradner JE, Kaelin WG, Jr. (2014) The myeloma drug lenalidomide promotes the cereblon-dependent destruction of Ikaros proteins. Science 343: 305–9

Maiques-Diaz A, Spencer GJ, Lynch JT, Ciceri F, Williams EL, Amaral FMR, Wiseman DH, Harris WJ, Li Y, Sahoo S, Hitchin JR, Mould DP, Fairweather EE, Waszkowycz B, Jordan AM, Smith DL, Somervaille TCP (2018) Enhancer Activation by Pharmacologic Displacement of LSD1 from GFI1 Induces Differentiation in Acute Myeloid Leukemia. Cell Rep 22: 3641–3659

Marcais A, Jeannet R, Hernandez L, Soulier J, Sigaux F, Chan S, Kastner P (2010) Genetic inactivation of Ikaros is a rare event in human T-ALL. Leuk Res 34: 426–9

McCarty AS, Kleiger G, Eisenberg D, Smale ST (2003) Selective dimerization of a C2H2 zinc finger subfamily. Mol Cell 11: 459–70

McClellan D, Casey MJ, Bareyan D, Lucente H, Ours C, Velinder M, Singer J, Lone MD, Sun W, Coria Y, Mason CC, Engel ME (2019) Growth Factor Independence 1B-Mediated Transcriptional Repression and Lineage Allocation Require Lysine-Specific Demethylase 1-Dependent Recruitment of the BHC Complex. Mol Cell Biol 39

Molnar A, Wu P, Largespada DA, Vortkamp A, Scherer S, Copeland NG, Jenkins NA, Bruns G, Georgopoulos K (1996) The Ikaros gene encodes a family of lymphocyte-restricted zinc finger DNA binding proteins, highly conserved in human and mouse. J Immunol 156: 585–92

Ngoenkam J, Schamel WW, Pongcharoen S (2018) Selected signalling proteins recruited to the T-cell receptor-CD3 complex. Immunology 153: 42–50

O’Neil J, Grim J, Strack P, Rao S, Tibbitts D, Winter C, Hardwick J, Welcker M, Meijerink JP, Pieters R, Draetta G, Sears R, Clurman BE, Look AT (2007) FBW7 mutations in leukemic cells mediate NOTCH pathway activation and resistance to gamma-secretase inhibitors. J Exp Med 204: 1813–24

Olsson A, Venkatasubramanian M, Chaudhri VK, Aronow BJ, Salomonis N, Singh H, Grimes HL (2016) Single-cell analysis of mixed-lineage states leading to a binary cell fate choice. Nature 537: 698–702

Person RE, Li FQ, Duan Z, Benson KF, Wechsler J, Papadaki HA, Eliopoulos G, Kaufman C, Bertolone SJ, Nakamoto B, Papayannopoulou T, Grimes HL, Horwitz M (2003) Mutations in proto-oncogene GFI1 cause human neutropenia and target ELA2. Nat Genet 34: 308–12

Peterson CL, Herskowitz I (1992) Characterization of the yeast SWI1, SWI2, and SWI3 genes, which encode a global activator of transcription. Cell 68: 573–83

Quentmeier H, Pommerenke C, Dirks WG, Eberth S, Koeppel M, MacLeod RAF, Nagel S, Steube K, Uphoff CC, Drexler HG (2019) The LL-100 panel: 100 cell lines for blood cancer studies. Sci Rep 9: 8218

Reynolds TC, Smith SD, Sklar J (1987) Analysis of DNA surrounding the breakpoints of chromosomal translocations involving the beta T cell receptor gene in human lymphoblastic neoplasms. Cell 50: 107–17

Rothenberg EV, Taghon T (2005) Molecular genetics of T cell development. Annu Rev Immunol 23: 601–49

Roux KJ, Kim DI, Raida M, Burke B (2012) A promiscuous biotin ligase fusion protein identifies proximal and interacting proteins in mammalian cells. J Cell Biol 196: 801–10

Saleque S, Kim J, Rooke HM, Orkin SH (2007) Epigenetic regulation of hematopoietic differentiation by Gfi-1 and Gfi-1b is mediated by the cofactors CoREST and LSD1. Mol Cell 27: 562–72

Shi LZ, Kalupahana NS, Turnis ME, Neale G, Hock H, Vignali DA, Chi H (2013) Inhibitory role of the transcription repressor Gfi1 in the generation of thymus-derived regulatory T cells. Proc Natl Acad Sci U S A 110: E3198–205

Sun L, Crotty ML, Sensel M, Sather H, Navara C, Nachman J, Steinherz PG, Gaynon PS, Seibel N, Mao C, Vassilev A, Reaman GH, Uckun FM (1999) Expression of dominant-negative Ikaros isoforms in T-cell acute lymphoblastic leukemia. Clin Cancer Res 5: 2112–20

Szklarczyk D, Gable AL, Lyon D, Junge A, Wyder S, Huerta-Cepas J, Simonovic M, Doncheva NT, Morris JH, Bork P, Jensen LJ, Mering CV (2019) STRING v11: protein-protein association networks with increased coverage, supporting functional discovery in genome-wide experimental datasets. Nucleic Acids Res 47: D607–D613

Terwilliger T, Abdul-Hay M (2017) Acute lymphoblastic leukemia: a comprehensive review and 2017 update. Blood Cancer J 7: e577

Tinsley KW, Hong C, Luckey MA, Park JY, Kim GY, Yoon HW, Keller HR, Sacks AJ, Feigenbaum L, Park JH (2013) Ikaros is required to survive positive selection and to maintain clonal diversity during T-cell development in the thymus. Blood 122: 2358–68

Tottone L, Zhdanovskaya N, Carmona Pestana A, Zampieri M, Simeoni F, Lazzari S, Ruocco V, Pelullo M, Caiafa P, Felli MP, Checquolo S, Bellavia D, Talora C, Screpanti I, Palermo R (2019) Histone Modifications Drive Aberrant Notch3 Expression/Activity and Growth in T-ALL. Front Oncol 9: 198

van der Meer LT, Jansen JH, van der Reijden BA (2010) Gfi1 and Gfi1b: key regulators of hematopoiesis. Leukemia 24: 1834–43

Velinder M, Singer J, Bareyan D, Meznarich J, Tracy CM, Fulcher JM, McClellan D, Lucente H, Franklin S, Sharma S, Engel ME (2017) GFI1 functions in transcriptional control and cell fate determination require SNAG domain methylation to recruit LSD1. Biochem J 474: 2951

Velu CS, Chaubey A, Phelan JD, Horman SR, Wunderlich M, Guzman ML, Jegga AG, Zeleznik-Le NJ, Chen J, Mulloy JC, Cancelas JA, Jordan CT, Aronow BJ, Marcucci G, Bhat B, Gebelein B, Grimes HL (2014) Therapeutic antagonists of microRNAs deplete leukemia-initiating cell activity. J Clin Invest 124: 222–36

Volpe G, Walton DS, Grainger DE, Ward C, Cauchy P, Blakemore D, Coleman DJL, Cockerill PN, Garcia P, Frampton J (2017) Prognostic significance of high GFI1 expression in AML of normal karyotype and its association with a FLT3-ITD signature. Sci Rep 7: 11148

Waegemans E, Van de Walle I, De Medts J, De Smedt M, Kerre T, Vandekerckhove B, Leclercq G, Wang T, Plum J, Taghon T (2014) Notch3 activation is sufficient but not required for inducing human T-lineage specification. J Immunol 193: 5997–6004

Winandy S, Wu P, Georgopoulos K (1995) A dominant mutation in the Ikaros gene leads to rapid development of leukemia and lymphoma. Cell 83: 289–99

Xu X, Choi SH, Hu T, Tiyanont K, Habets R, Groot AJ, Vooijs M, Aster JC, Chopra R, Fryer C, Blacklow SC (2015) Insights into Autoregulation of Notch3 from Structural and Functional Studies of Its Negative Regulatory Region. Structure 23: 1227–35

Yucel R, Karsunky H, Klein-Hitpass L, Moroy T (2003) The transcriptional repressor Gfi1 affects development of early, uncommitted c-Kit+ T cell progenitors and CD4/CD8 lineage decision in the thymus. J Exp Med 197: 831–44

Zarebski A, Velu CS, Baktula AM, Bourdeau T, Horman SR, Basu S, Bertolone SJ, Horwitz M, Hildeman DA, Trent JO, Grimes HL (2008) Mutations in growth factor independent-1 associated with human neutropenia block murine granulopoiesis through colony stimulating factor-1. Immunity 28: 370–80

Zeng H, Yucel R, Kosan C, Klein-Hitpass L, Moroy T (2004) Transcription factor Gfi1 regulates self-renewal and engraftment of hematopoietic stem cells. EMBO J 23: 4116–25

Zhang J, Ding L, Holmfeldt L, Wu G, Heatley SL, Payne-Turner D, Easton J, Chen X, Wang J, Rusch M, Lu C, Chen SC, Wei L, Collins-Underwood JR, Ma J, Roberts KG, Pounds SB, Ulyanov A, Becksfort J, Gupta P et al. (2012) The genetic basis of early T-cell precursor acute lymphoblastic leukaemia. Nature 481: 157–63

Zhang J, Jackson AF, Naito T, Dose M, Seavitt J, Liu F, Heller EJ, Kashiwagi M, Yoshida T, Gounari F, Petrie HT, Georgopoulos K (2011) Harnessing of the nucleosome-remodeling-deacetylase complex controls lymphocyte development and prevents leukemogenesis. Nat Immunol 13: 86–94

